# Microenvironment Shapes Small Cell Lung Cancer Neuroendocrine States and Presents Therapeutic Opportunities

**DOI:** 10.1101/2024.02.09.579028

**Authors:** Parth Desai, Nobuyuki Takahashi, Rajesh Kumar, Samantha Nichols, Justin Malin, Allison Hunt, Christopher Schultz, Yingying Cao, Desiree Tillo, Darryl Nousome, Lakshya Chauhan, Linda Sciuto, Kimberly Jordan, Vinodh Rajapakse, Mayank Tandon, Delphine Lissa, Yang Zhang, Suresh Kumar, Lorinc Pongor, Abhay Singh, Brett Schroder, Ajit Kumar Sharma, Tiangen Chang, Rasa Vilimas, Danielle Pinkiert, Chante Graham, Donna Butcher, Andrew Warner, Robin Sebastian, Mimi Mahon, Karen Baker, Jennifer Cheng, Ann Berger, Ross Lake, Melissa Abel, Manan Krishnamurthy, George Chrisafis, Peter Fitzgerald, Micheal Nirula, Shubhank Goyal, Devon Atkinson, Nicholas W. Bateman, Tamara Abulez, Govind Nair, Andrea Apolo, Udayan Guha, Baktiar Karim, Rajaa El Meskini, Zoe Weaver Ohler, Mohit Kumar Jolly, Alejandro Schaffer, Eytan Ruppin, David Kleiner, Markku Miettinen, G Tom Brown, Stephen Hewitt, Thomas Conrads, Anish Thomas

**Affiliations:** Developmental Therapeutics Branch, Center for Cancer Research, National Cancer Institute, National Institutes of Health, Bethesda, Maryland, USA; Department of Medical Oncology, Fox Chase Cancer Center, Temple University Hospital and Lewis Katz School of Medicine, Philadelphia, Pennsylvania; Department of Medical Oncology, National Cancer Center Hospital East, Kashiwa, Japan; Women’s Health Integrated Research Center, Inova Health System, Falls Church, Virginia, USA; Cancer Data Science Laboratory, Center for Cancer Research, National Cancer Institute, National Institutes of Health, Bethesda, Maryland, USA; CCR Collaborative Bioinformatics, Resource, Office of Science and Technology Resources, National Cancer Institute, National Institutes of Health, Bethesda, MD 20892, USA; Center for Biosystems Science and Engineering, Indian Institute of Science, Bangalore, India; Department of Immunology and Microbiology, University of Colorado Anschutz Medical Campus, Aurora, Colorado, USA; Laboratory of Human Carcinogenesis, Center for Cancer Research National Cancer Institute, National Institutes of Health, Bethesda, Maryland, USA; HCEMM Cancer Genomics and Epigenetics Research Group, Szeged 6728, Hungary; Molecular Histopathology Laboratory, Laboratory Animal Sciences Program, Frederick National Laboratory for Cancer Research, National Cancer Institute, National Institutes of Health, Frederick, Maryland, USA; Pain and Palliative care services, National Institutes of Health Clinical Center, Bethesda, Maryland, USA; Laboratory of Genitourinary cancer Pathogenesis, Center for Cancer Research, National Cancer Institute, National Institutes of Health, Bethesda, Maryland, USA; Center for Advanced Preclinical Research, Frederick National Laboratory for Cancer Research, National Cancer Institute, National Institutes of Health, Frederick, Maryland, USA; The Henry M. Jackson Foundation for the Advancement of Military Medicine Inc., Bethesda, MD, 20817, USA; National Institute of Neurological Disorders and Stroke, Center for Cancer Research, National Cancer Institute, National Institutes of Health, Bethesda, Maryland, USA; Genitourinary Malignancies Branch, Center for Cancer Research, National Cancer Institute, National Institutes of Health, Bethesda, Maryland, USA; Thoracic and GI Malignancies Branch, Center for Cancer Research, National Cancer Institute, National Institutes of Health, Bethesda, Maryland, USA; Laboratory of Pathology, National Cancer Institute, National Institutes of Health, Bethesda, Maryland, USA

**Keywords:** small cell lung cancer, tumor microenvironment, tumor heterogeneity, spatial transcriptomics, cancer associated fibroblasts, intercellular communication, rapid research autopsy

## Abstract

Small-cell lung cancer (SCLC) is the most fatal form of lung cancer. Intra-tumoral heterogeneity, marked by neuroendocrine (NE) and non-neuroendocrine (non-NE) cell states, defines SCLC, but the drivers of SCLC plasticity are poorly understood. To map the landscape of SCLC tumor microenvironment (TME), we apply spatially resolved transcriptomics and quantitative mass spectrometry-based proteomics to metastatic SCLC tumors obtained via rapid autopsy. The phenotype and overall composition of non-malignant cells in the tumor microenvironment (TME) exhibits substantial variability, closely mirroring the tumor phenotype, suggesting TME-driven reprogramming of NE cell states. We identify cancer-associated fibroblasts (CAF) as a crucial element of SCLC TME heterogeneity, contributing to immune exclusion, and predicting exceptionally poor prognosis. Together, our work provides a comprehensive map of SCLC tumor and TME ecosystems, emphasizing their pivotal role in SCLCs adaptable nature, opening possibilities for re-programming the intercellular communications that shape SCLC tumor states.

## Introduction

Intratumor heterogeneity is a fundamental problem in cancer^1^. A major contributor to intratumor heterogeneity is phenotypic plasticity, which endows tumor cells with the ability to assume distinct cell identities, enabling metastatic capabilities and drug resistance^2,3^. A better understanding of the determinants of phenotypic plasticity is critical and may aid manipulation of cancer cell states and targeting the associated vulnerabilities.

Small-cell lung cancer (SCLC), a high-grade neuroendocrine cancer, represents a paradigm to study tumor heterogeneity and its consequences. As the most metastatic, treatment-resistant, and fatal form of lung cancer^4,5^, SCLC exhibits a high degree of intratumoral heterogeneity, harboring cells of neuroendocrine (NE) and non-neuroendocrine (non-NE) states^6–12^, further defined by differential expression of lineage-defining transcription factors *ASCL1, NEUROD1*, and *POU2F3*. A fourth subtype has been characterized by *YAP1* expression^9,11,13–15^ or low expression of all three transcription factors accompanied by an inflamed gene expression program^16^. SCLC subtypes, defined by the dominant cell states in each tumor, exhibit distinct therapeutic vulnerabilities^10,11^. Immunogenic plasticity and Notch signaling of non-NE SCLC underlie their favorable responses to immune checkpoint blockade^17^. NE SCLC is characterized by replication stress, rendering them susceptible to DNA repair-targeted agents^10,18–22^. While heterogeneity and plasticity are important determinants of SCLC clinical responses, the origins and organization of SCLC heterogeneity are poorly understood^12,16,23,24^.

Tumor cell state represents the combined influences of both cell-intrinsic (e.g., mutational background, epigenetic state) and cell-extrinsic (e.g., cell-to-cell interactions, microenvironment) factors^25^. In genetically engineered mouse models (GEMMs), the cell-of-origin profoundly influences the SCLC cell states^26,27^. Notch signaling, generally suppressed in NE SCLC, can induce a transition from NE to non-NE cell state^28,29^. MYC, frequently amplified on extrachromosomal DNA^30^, can activate Notch signaling to promote the temporal evolution of SCLC sequentially from an ASCL1 to a NEUROD1 to a non-NE state^28,31^. However, model systems that have informed SCLC biology to date harbor minimal to no tumor microenvironment (TME)^27,32^, and as such, very little is known of the cell-extrinsic drivers of SCLC tumor state. Importantly, SCLC TME has not been examined in depth in human tumors, especially in a spatial context, owing to the challenges of obtaining tumor samples. SCLC is often diagnosed using fine needle aspirates, and biopsies at relapse are not standard. Research biopsies are difficult to obtain due to rapid cancer progression and patient comorbidities, and when available, they may not portray the true extent of TME heterogeneity^33^. Further, the sequencing approaches that have been applied to human SCLC to date, including bulk and single-cell RNA-sequencing (scRNA-seq), do not inform the spatial interactions between tumor cells and TME.

Thus, while most patients with SCLC are diagnosed with and succumb to metastatic disease^34^, the current understanding of SCLC heterogeneity, derived largely from model systems, is limited to cell-intrinsic influences^27^. Our understanding of cell-extrinsic influences on SCLC plasticity is rudimentary, but a key role of these non-genetic mechanisms is suggested by two important observations: (i) transcriptional subtypes of human SCLC are not strongly associated with specific mutational patterns^35^ and (ii) divergence of NE gene expression programs between human tumors and patient-derived xenografts which lack human TME^9^. We hypothesized that dynamic interactions between TME and tumor cells shape SCLC tumor states, plasticity, and heterogeneity. Here, we studied metastatic and treatment-resistant SCLC in patients who underwent research autopsies and applied spatially resolved transcriptomics, integrating the data with whole-genome sequencing, bulk, and immunohistochemistry (IHC), multispectral imaging of multiplex immunofluorescence, and mass spectrometry (MS)-based proteomics seeking to map the cell-extrinsic factors that shape SCLC cell states in their positional context.

## Results

### Patient and tumor characteristics

SCLC surgical resections are rarely performed since the tumors are almost always widely metastatic by the time of diagnosis^34^. Diagnostic core needle biopsies and cytology samples provide an inadequate representation of the TME heterogeneity^33^. Thus, to characterize human SCLC tumor and TME in its spatial context, we performed rapid research autopsies on ten patients (Clinicaltrials.gov identifier: NCT01851395) and profiled formalin-fixed, paraffin-embedded (FFPE) tumors using NanoString GeoMx Digital Spatial Profiler and GeoMx Whole Transcriptome Atlas (Methods; **Fig. 1A**, Fig. S1A). Patients were mostly male and smokers (8 of 10 each), with a median age of 64 (range: 47-75 years) (Table S1). All patients had received at least two systemic therapies including platinum and etoposide, and immunotherapy in eight of ten cases. The study included a single metastatic site from each patient, selected to represent frequent sites of SCLC metastases^36^; seven derived from liver, two lymph nodes, and one adrenal. Expert pathologists reviewed tumor sections to confirm the diagnoses. Nine out of 10 tumors were morphologically classified as SCLC. One tumor had combined morphology, with SCLC and squamous differentiation (patient #7). Patient #6 had a diagnostic biopsy of combined small cell carcinoma with adenocarcinoma differentiation, but only small cell morphology was observed on the autopsy tumor. All tumors underwent whole genome sequencing (WGS) and six of ten tumors were also laser micro-dissected and profiled using mass spectrometry-based proteomics. Mutations or copy number alterations of *TP53* and *RB1* were observed in most (*RB1* in 7 of 10 and *TP53* in 6 of 10) tumors (Fig. S1B). Genomic events that functionally mimic *TP53* and *RB1* inactivation^5^, such as *MDM2* or *CCND1* amplifications were found in three tumors. *MYC* family genes were amplified (>4 copy numbers) in six cases. The two never-smoker tumors had no known pathogenic *TP53* alterations detected but had *MYCL* amplification and *RB1* deletion in one case and *CCND1* amplification in the other.

**Figure 1:**
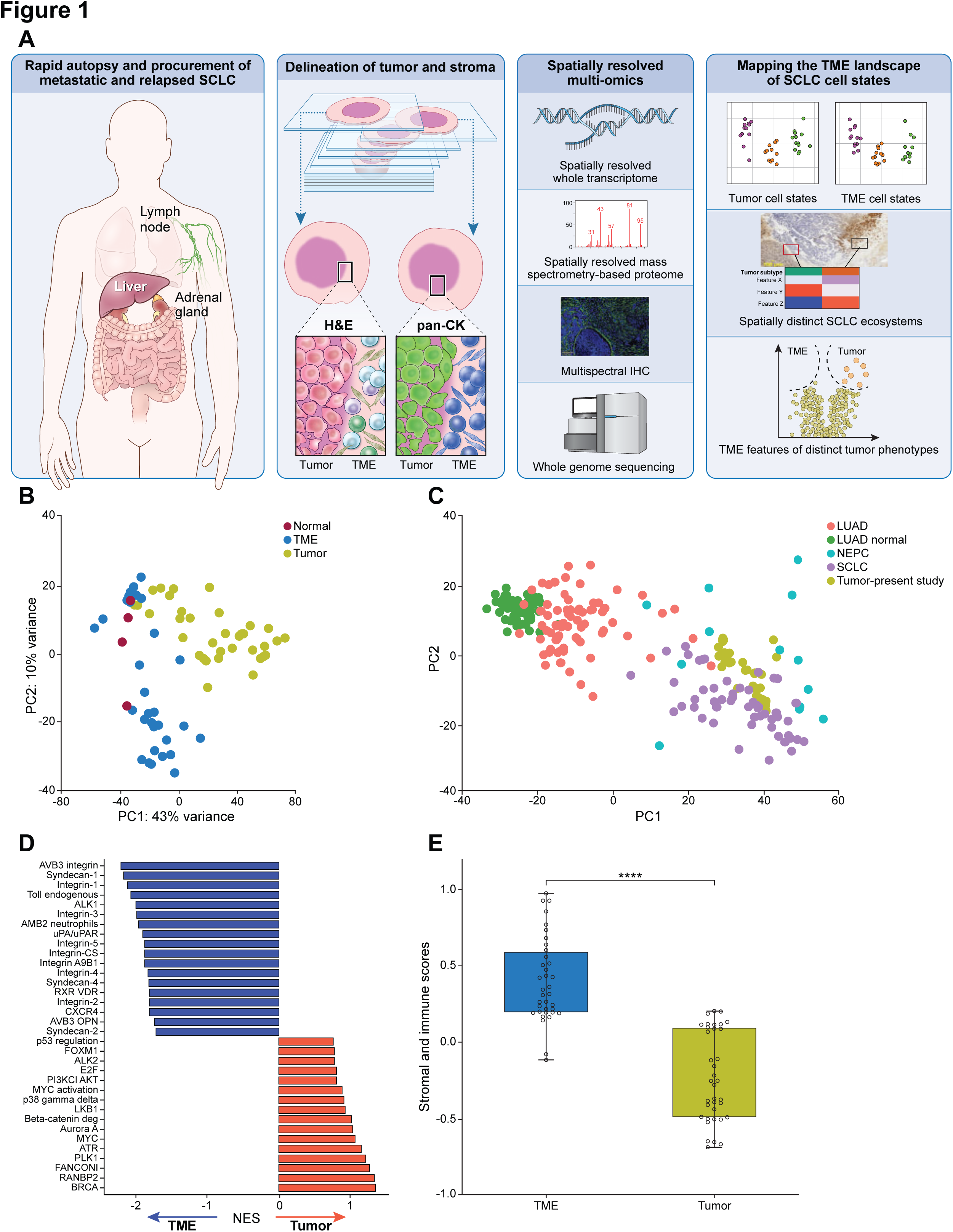
Dissection of metastatic and relapsed SCLC using spatial transcriptomics. A. Workflow of SCLC tumor sampling, tissue sectioning, genomics, and spatially resolved transcriptomics and proteomics. B. PCA of gene expression derived from tumor (n=36), TME (n=30), and normal (n=4) segments, 2500 genes with highest variance. C. Projection of tumor segments (n=36) to PCA performed on lung adenocarcinoma, NEPC, SCLC, and adjacent normal lung gene expression^38^. D. NES (GSEA) of differentially expressed PID pathways^90^ between tumor and TME segments. E. Stromal and immune score^40^ (ssGSEA) computed for TME and tumor segments^#^ Abbreviations: Pan CK, pan-cytokeratin; SCLC, small cell lung cancer; NEPC, neuroendocrine prostate cancer; LUAD, Lung adenocarcinoma; LUAD normal, adjacent normal lung; NES, normalized enrichment score; TME, tumor microenvironment; GSEA, gene set enrichment analysis; ssGSEA, single sample gene set enrichment analysis; PID, pathway interaction database; **** statistical significance at p<0.0001; ^#^ student t-test.

Tissue sections were profiled using fluorescently labeled antibodies targeting epithelial cells (pan-cytokeratin) and immune cells (CD45) to discriminate tumor and TME in selected regions, measuring an average 600-micron diameter (range, 400-650), with an average of three regions (range, 3-6) per tumor section. Tumor and TME were profiled independently based on pan-cytokeratin staining and morphological assessment, even when the two compartments were adjoined or interdigitated, yielding two whole transcriptome profiles per region of interest (see extended data). Four tumor-only areas with minimal visible TME and four histologically normal areas with no visible surrounding tumor were also profiled. Thus, transcriptomes of 72 regions, including 36 tumor, 32 TME, and four normal segments were generated from ten tumors. Barcoded oligonucleotide probes were designed for a total of 18,677 genes representing the whole transcriptome^37^. Following probe hybridization, ultraviolet cleavage, and barcode collection, gene expression was quantified by polymerase chain reaction (PCR) amplification and Illumina sequencing. Two TME segments that did not meet the sequencing quality metrics (Fig. S1C) were excluded from further analyses. While TME and tumor segments had comparable areas of capture, more nuclei were profiled from tumor segments than TME segments (Fig. S1D). Sequencing saturation was high (>90%) for all segments. Sublevel sections of areas profiled for gene expression were additionally examined using IHC, multiplex protein immunofluorescence, whole genome sequencing, and mass spectrometry-based proteomics.

### Distinct gene expression profiles of SCLC tumor and TME

To characterize the distinct transcriptomic features of tumor and TME, we evaluated the top 2500 variably expressed genes across all segments (**Fig. 1B**). This analysis revealed separate clustering of tumor and TME. Maximum variance was observed in the principal component (PC)1 axis which captured 43% of differences between tumor and TME. The histologically normal-appearing tissue adjacent to tumor clustered together with TME underscoring similarities between them. However, at a patient level, the normal tissue clustered separately from tumor and TME (Fig. S1E). Tumor gene expression profiles were similar to that of SCLC tumors^5^ and neuroendocrine prostate cancer (NEPC)^38^ (**Fig. 1C**) and distinct from lung adenocarcinoma and normal lung.

We applied several analytical approaches to better understand the specificity of tumor and TME gene expression profiles. Segments annotated as tumor exhibited higher NE gene expression scores^9,39^ than TME and normal segments (Fig. S1F), consistent with the characteristic expression of NE genes in SCLCs. Gene Set Enrichment Analysis (GSEA) showed upregulation of pathways related to DNA repair, replication stress, and MYC in tumors, compared with TME which were enriched for pathways related to extracellular matrix and inflammation (**Fig. 1D**). Single sample Gene Set Enrichment Analysis (ssGSEA)^40^ demonstrated significantly higher inferred fraction of stroma and immune cells in TME compared with tumors (**Fig. 1E**). SCLC tumors with lower NE scores^39^ had higher stromal and immune scores (Fig. S1G), consistent with intrinsic tumor immunity of non-NE SCLC^16,17^. Estimated tumor purity of most tumor segments were more than 95% and exceeded those of bulk-tumor derived estimates^5,9,41^ (Fig. S1H). Thus, despite its spatial proximity, SCLC tumor and TME harbor distinct gene expression states and programs.

### Spatial Intra-tumoral heterogeneity of SCLC neuroendocrine differentiation

Seeking to classify SCLC tumors in an unbiased manner, we identified the optimal number of clusters for the 36 tumor segments as k=3 (see methods; Fig. S2A). The three clusters partitioned clearly at 2500 highly variant genes (**Fig. 2A**). In four of ten tumors profiled (Patients #3, 5, 7, 10), spatially proximate segments of the same tumor were separated into different clusters (Fig. S2B). Cluster 1 showed relatively high expression of *Notch* genes and *REST*, a repressor of neural gene expression and direct target of *Notch1* (**Fig. 2B**, Fig. S2C). Consistent with the negative regulation of NE differentiation by Notch, cluster 1 showed reduced expression of several key NE genes (*INSM1*, *BEX1*, *NCAM1*). In contrast, Cluster 3 showed upregulation of NE genes and Notch inhibitory ligands *DLL1* and *DLL3*. Cluster 2 exhibited features of both clusters 1 and 3, including simultaneous expression of both NE and non-NE genes^7^. Additionally, they expressed multiple EMT genes (*FN1, SNAI2*), in contrast to clusters 1 and 3 which showed variable expression of EMT genes. Cluster 2 also showed co-expression of mesenchymal and epithelial genes *CD44* and *EPCAM* that are characteristically expressed in non-NE and NE cells respectively^42^. *MYC* was amplified in bulk genome sequences of cluster 1-containing tumors, consistent with the known role of MYC in driving a non-NE phenotype^14,28,31^. However, the tumor clusters were not defined by genomic alterations including mutations and or copy number alterations of *TP53* or *RB1* (Fig. S2D).

**Figure 2:**
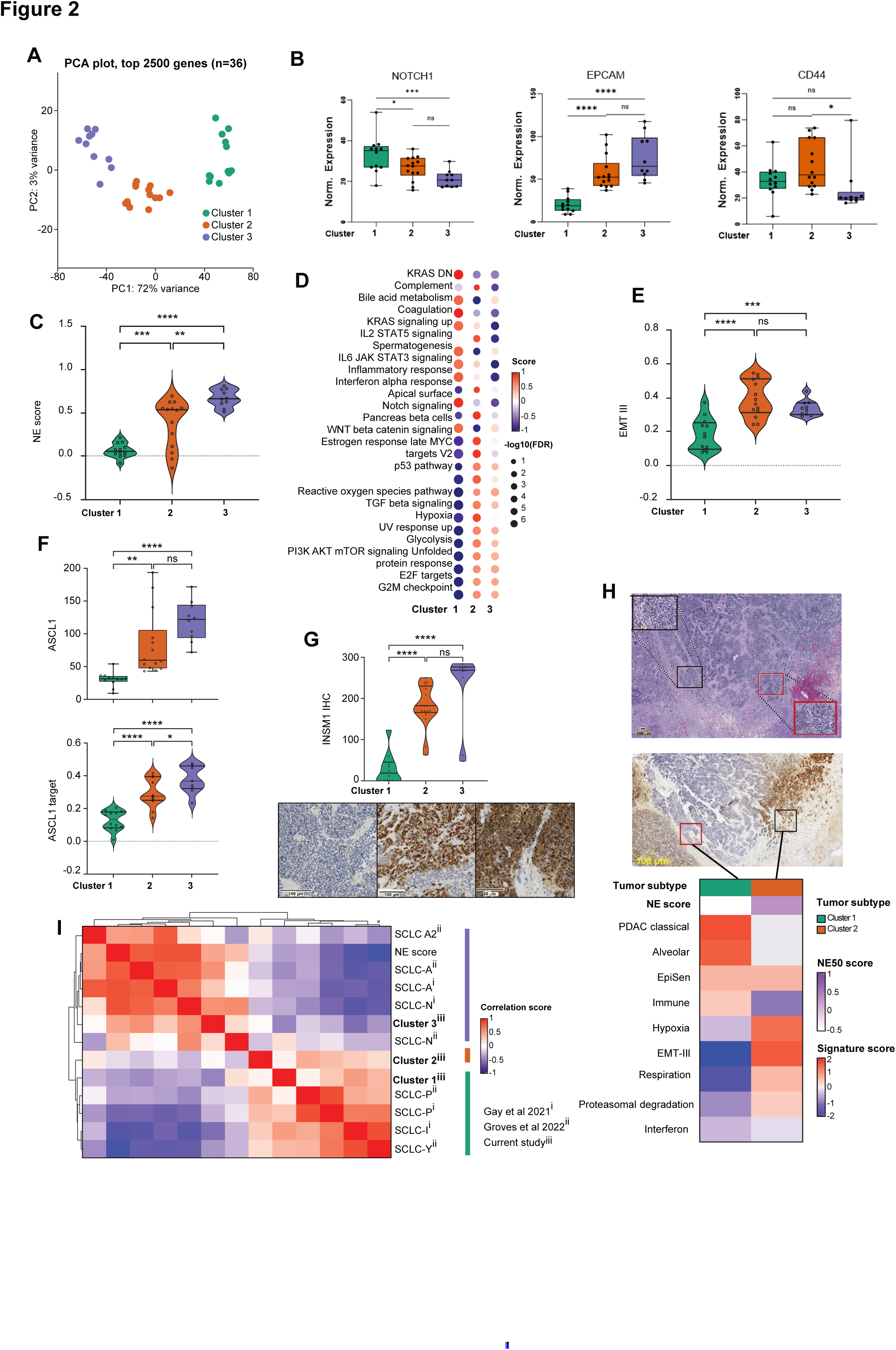
Spatial intra-tumoral heterogeneity of SCLC neuroendocrine differentiation. A. PCA of 2500 genes with highest variance across tumor segments (n=36). B. Normalized expression of NOTCH1, EPCAM and CD44 for the three tumor segment clusters^#^. C. NE scores across the tumor clusters^#^. D. Differentially enriched Hallmark pathways (GSEA) across the tumor clusters^$^. E. EMT III^44^ (ssGSEA) scores across the tumor clusters^#^. F. *ASCL1* expression (top, Q3 value) and *ASCL1* target scores (bottom, ssGSEA) across the three tumor segment clusters^#^. G. INSM1 protein expression across the tumor clusters. H-score range from 0-300^#^. H. Hybrid-NE (red square) and non-NE (black square) regions in morphologically similar and spatially proximate segments of the same tumor (patient #10). H/E, INSM1 IHC, and differentially upregulated cancer metaprograms^44^ shown. Medium power (20x), high power (40x) inset. Scale bar at 100 µm. I. Pairwise correlation of tumor-cluster signatures from current study and previously published SCLC gene signatures^7,16,51^ computed on SCLC tumor transcriptomes (n=81)^5^. Abbreviations: SCLC, small cell lung carcinoma; PCA, principal component analysis; ssGSEA, single sample gene set enrichment analysis; FDR, false discovery rate; EMT, Epithelial mesenchymal transformation; TF, transcription factors; ASCL1, Achaete-scute complex homologue 1; NEUROD1, neuronal differentiation 1; YAP1, yes-associated protein 1; POU2F3, POU class 2 homeobox 3; Q3, 3^rd^ quantile value; UMAP, uniform manifold approximation and projection; NE, neuroendocrine; INSM1, insulinoma-associated protein 1; NES, normalized enrichment score; IHC, Immunohistochemistry; KRAS DN, KRAS downregulation; ns, statistically non-significant; H/E-hematoxylin & Eosin; *statistical significance at p<0.05; **statistical significance at p<0.001; ***statistical significance at p<0.001; ****statistical significance at p<0.0001; ^#^Tukey’s multiple comparison test; ^$^FDR correction using Benjamini and Hochberg (BH) method.

In line with these observations at the gene expression level, cluster 1 exhibited significantly lower NE gene signature score^39^ (mean=0.06; range, –0.07 to 0.21) than cluster 3 which showed the highest NE score (mean=0.67, range=0.51 to 0.81), and cluster 2 (mean=0.38; range, –0.13 to 0.69) which had intermediate NE scores (**Fig. 2C**). Consistently, cluster 3 was enriched for NE SCLC hallmarks including DNA repair and replication stress^18^, E2F targets, and G2M checkpoints^10^ (**Fig. 2D**, Fig. S2E). Cluster 1 was enriched for non-NE SCLC hallmarks such as inflammation and immunity^9,43^. Cluster 2 shared features of both clusters 1 and 3, including replication stress, E2F targets, G2M checkpoint, and immune pathways. Additionally, cluster 2 exhibited selective upregulation of hypoxia and epithelial-mesenchymal transition (EMT) III^44^ (**Fig. 2E**). EMT-III is a cancer metaprogram of coordinately upregulated mesenchymal and epithelial markers consistent with a hybrid cellular state, previously described in neuroendocrine cancers^44^.

Transcription factors function as molecular switches to regulate the expression of cell-type or lineage-specific target genes. Clusters 2 and 3 exhibited significantly higher expression of NE lineage-defining transcription factor *ASCL1* and its downstream target genes (**Fig. 2F**) than *NEUROD1* or non-NE lineage defining *POU2F3* (Fig. S2F). Cluster 1 exhibited low and comparable expression of all four transcription factors and their downstream target genes, reminiscent of SCLC-I (inflamed subtype) (Fig. S2G)^16^. IHC of sublevel sections (Table S2) confirmed these observations. The highest ASCL1 protein expression was observed in cluster 3, followed by cluster 2, and minimal expression in cluster 1. NEUROD1 and POU2F3 were rarely expressed across the three clusters (Fig. S2H). INSM1, a transcription factor that regulates global neuroendocrine gene programs^45^ was also significantly highly expressed in clusters 3 and 2 compared with cluster 1 (**Fig. 2G**). YAP1 protein was expressed in cluster 2, but importantly these tumors did not express *YAP1* transcriptional target signature (Fig. S2I) or a pan-cancer *YAP1*/*HIPPO* signature^46^ (Fig. S2J). In contrast, YAP1 protein was highly expressed in TME segments of cluster 2, and to a lesser extent cluster 3, but with concomitant upregulation of *YAP1* target genes in the TME. The functional role of stromal *YAP1* signaling in cluster 2 and 3 remains to be explored in future studies.

Morphologically all three SCLC clusters exhibited similar nuclear and cytoplasmic features. However, Cluster 2 tumors were more likely to localize at the invasive margin of tumors, forming nests and buds surrounded by desmoplastic tissue^47^ (Fig. S2K). In contrast, Cluster 1 and Cluster 3 tumors formed large sheets or closely packed interconnected ribbons. GSEA of cancer hallmark capabilities^48,49^ revealed striking enrichment of nearly every cancer hallmark in Cluster 2 tumors (Fig. S2L) compared with the other two subtypes indicating distinctly aggressive features of this phenotype. Even within the same tumor specimen, despite spatial proximity and morphological similarity between the unique segments, Cluster 2 tumor segments exhibited (**Fig. 2H**, S2M, Table S3) specific upregulation of EMT-III^44^ and hypoxia pathways. Given the technical constraints of our approach, which does not achieve single-cell resolution, we explored the potential for Cluster 2 tumors to represent potential contamination of tumor and stromal cells. The tumor and TME components of Cluster 2 segments distinctly separated along PC1 versus PC2 (Fig. S2N), and the Cluster 2 tumor segments demonstrated significantly lower stromal and immune scores compared to the corresponding Cluster 2 TME segments (Fig. S2O). Although, we cannot fully exclude the possibility that Cluster 2 segments represent a mixture of NE or non-NE cells, the observation of this phenotype is reminiscent of transitional states identified in single cell studies^50^ and in-vitro identification of cells with intermediate activation of Notch pathway^29^.

To benchmark cluster 2 and to clarify potential inter-relationships, we assessed how features of cluster 2 overlapped with previously described SCLC phenotypes^7,16,51^. We created cluster-specific gene signatures from the top contributors to the first and second PCs (Methods, Table S4). Pairwise correlations of these signatures revealed strong correlation of cluster 1 with the non-NE SCLC and cluster 3 with NE SCLC signatures. However, cluster 2 was not well described by prior signatures (**Fig. 2I**, S2P). Tumor clusters 1, 2 and 3 are henceforth referred to as non-NE, hybrid-NE, and NE tumor subtypes respectively. This classification is supported by the relative expression of canonical transcription factors, their transcriptional targets, gene expression programs, and matched protein expression.

### Tumor heterogeneity-linked reprogramming of SCLC TME

We next examined the TME in relation to the spatially proximate tumor subtypes. PCA of 5000 most variable genes of the 30 TME segments revealed clear separation to three TME clusters (**Fig. 3A**). NE and hybrid-NE TME were more like each other than non-NE TME.

**Figure 3:**
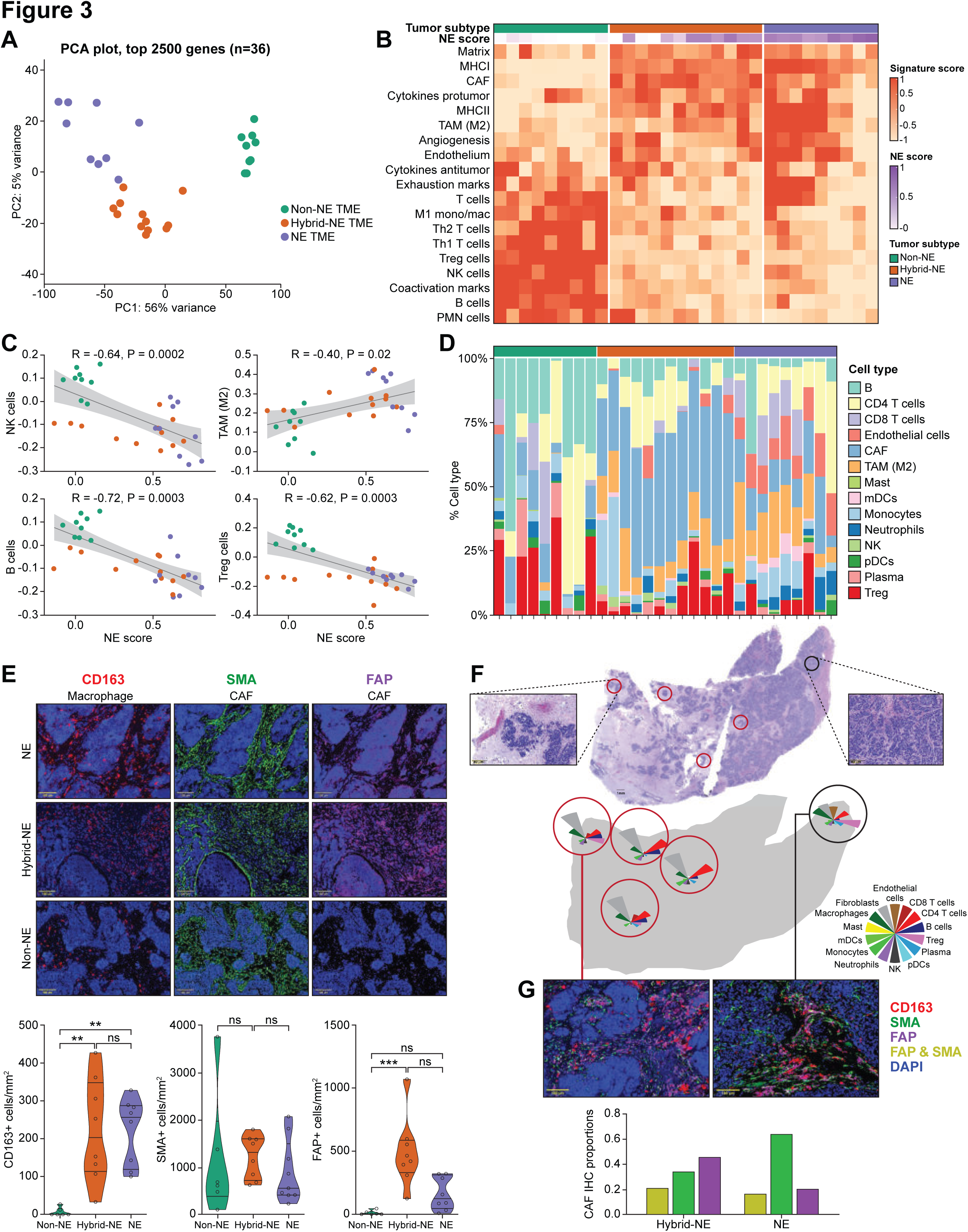
Tumor heterogeneity-linked reprogramming of SCLC TME. A. PCA of 5000 genes with highest variance across the TME segments (n=30). B. Pan-cancer TME signatures enriched across the TME subtype segments (n=30)^52^. C. Correlation between tumor segment NE scores and immune cell (NK-cell, TAM, B-cell, T-reg) signatures of the spatially proximate TME segment. NE subtypes are indicated in colors (color code as Fig 3A). D. Relative proportion of TME non-malignant cell types sorted by NE subtype of the spatially proximate tumor segments (n=30), estimated using CIBERSORT^57^. E. Representative single component multiplex immunofluorescence images showing CD163 (left), SMA (center) and FAP (right) expression across TME subtypes (n=30). Quantification shown below cells/mm^2#^. DAPI filter (nuclear stain) is applied on all the images. Scale bar 100 µm. F. Heterogeneity across spatially proximate hybrid-NE (n=4) (red circles) and NE (n=1) (black circle) TME within tumor from patient #5. Top panel shows bird’s eye view (4x magnification) H&E image of the tumor section with insets highlighting hybrid-NE (left) and NE (right) regions. Bottom panel shows CIBERSORT-derived relative TME cell type abundance^53^. Inset scale bar 100 µm. G. Enrichment of FAP+ cells in hybrid-NE TME. Representative (40x magnification) multispectral mIF images of hybrid-NE (left) and NE (right) TME from patient #5 in Fig 3F. Bar plot (below) demonstrating proportion of FAP+ cells, SMA + cells, and combined FAP and SMA+ cells in hybrid-NE (left) and NE (right) TME regions. DAPI (nuclear) filter is on in all the images. Abbreviations: PCA, principal component analysis; TME, tumor microenvironment; NE, neuroendocrine; mIF, multiplex immunofluorescence; ns, non-significant; DAPI, 41,6-diamidino-2-phenylindole; NE, neuroendocrine; FAP, fibroblast activation protein; SMA, smooth muscle actin; TAM, tumor associated macrophages; T-reg, regulatory T-cells; pDC; plasmacytoid dendritic cells; mDC, myeloid dendritic cells; H&E, Hematoxylin and eosin; *statistical significance at p<0.05; **statistical significance at p<0.01; ***statistical significance at p<0.001; R= spearman’s correlation co-efficient; ^#^Tukey’s multiple comparison test.

Applying pan-cancer TME signatures^52^, non-NE TME were marked by immune infiltration including natural killer (NK) cells, B cells, and M1 tumor associated macrophages (TAM-M1), and upregulation of immune co-activators (*CD40LG, CD80*) (**Fig. 3B**) compared with hybrid-NE and NE TME^9,43^. Non-NE TME were also enriched for regulatory T-cells (T-regs), immune checkpoints (*PDCD1, LAG3, TIGIT*), and neutrophils. In contrast, hybrid-NE and NE TME were characterized by distinct signatures including cancer associated fibroblasts (CAF) and TAM-M2, matrix remodeling, and pro-tumoral cytokines. The composition of individual immune cell types was correlated with NE differentiation (TAM-M2, r= 0.40, p=0.02; NK cells, r= –0.64, p=0.0002; B cells, r= –0.72, p=0.0003; T-regs, r= –0.52, p=0.0003) (**Fig. 3C**), suggesting reprogramming of the TME and individual immune cell types linked to tumor NE differentiation. Consistent observations were noted when the intratumoral proportion of immune cells was estimated based on CIBERSORT deconvolution^53^ (**Fig. 3D**), but additionally revealed a striking enrichment of immunosuppressive CAFs in hybrid-NE TME. Consistent with CAFs constituting the dominant cell-type in hybrid-NE TME, we found significantly lower Shannon index scores^54^ in these TMEs (Fig. S3A), especially compared to NE TME (p=0.0004). Additionally, hybrid-NE TME showed conspicuous absence of CD8+ T cell signatures (Fig. S3B).

To further validate these observations, we performed IHC and multiplex immunofluorescence on sublevel tumor sections to characterize B cells (CD20), T-cells (CD3), fibroblasts (SMA, FAP), macrophages (CD163, CD115, CD11b), and HLA-DR positive cells. Non-NE TME showed higher CD20+ B cell infiltrates compared with NE and hybrid-NE TME, consistent with transcriptomic data (Fig. S3C). Confirming CIBERSORT observations, non-NE and NE TME showed higher CD3+ T cell infiltrates compared with hybrid-NE TME which lacked CD3+ T cells corroborating the earlier observation of reduced intrinsic immune activation in hybrid-NE cells (Fig. 2I). NE and hybrid-NE TME were enriched for TAMs (CD163+ cells) (**Fig. 3E**, S3D) and CD115+ macrophages representing tumor infiltrating mono/macrophages compared to HLA-DR rich regulatory macrophages (Fig. S3E)^55^. All three TME subtypes showed similar proportions of SMA+ fibroblasts. However, hybrid-NE exhibited distinctly increased FAP+ fibroblast signals, while they were absent in non-NE TME.

Intra-tumoral heterogeneity between spatially separated TME regions also correlated with tumor NE subtypes (Fig. S3F). Intra-tumor TME heterogeneity was most evident in tumors that harbored at least one hybrid-NE region (patient #3, #10 and #5). For example, the TME compositions of the NE and hybrid-NE regions from liver metastasis of patient #5 were remarkably distinct (**Fig. 3F**). The hybrid-NE tumors showed budding and nesting, reduced CD8+T cells signatures and enrichment of CAF signatures. Multiplex immunofluorescence confirmed significant enrichment of FAP+ and FAP/SMA+ co-expressing cells in hybrid-NE TME (**Fig 3G**, S3G). Taken together, our observations suggest that NE differentiation is a major determinant of SCLC TME heterogeneity, favoring a highly active immune milieu in non-NE TME. The NE and hybrid-NE TME harbored more primitive immune cell profiles^56^, consistent with the known evolutionary trajectory of SCLC from NE to non-NE cell states. Notably, there was remarkable heterogeneity of CAF states across the tumor subtypes, with hybrid-NE subtype specifically enriched for FAP+ CAFs.

### Immunosuppressive CAF cell state enriched in hybrid-NE SCLC

We profiled cell states and multicellular communities that organize as functional units^57^, referred to as ecotypes, in the spatially resolved gene expression data (**Fig. 4A**, Table S5). NE and hybrid-NE subtypes were enriched for carcinoma ecotype 1 (CE1), associated with lymphocyte deficiency, EMT, and characteristic of cancers with the poorest prognosis^57^. NE and hybrid-NE subtypes additionally showed enrichment of CAF S3 cell state, marked by expression of extracellular matrix, and collagen organization and degradation-related pathways (**Fig. 4B**) and macrophage S4 (Mac S4) (Fig. S4A). In contrast, non-NE TME were enriched for CAF S6, characterized by pathways related to neuronal system, G-coupled protein mediated receptor ligand binding, and peptide ligand binding receptor activity (Fig. S4B).

**Figure 4.**
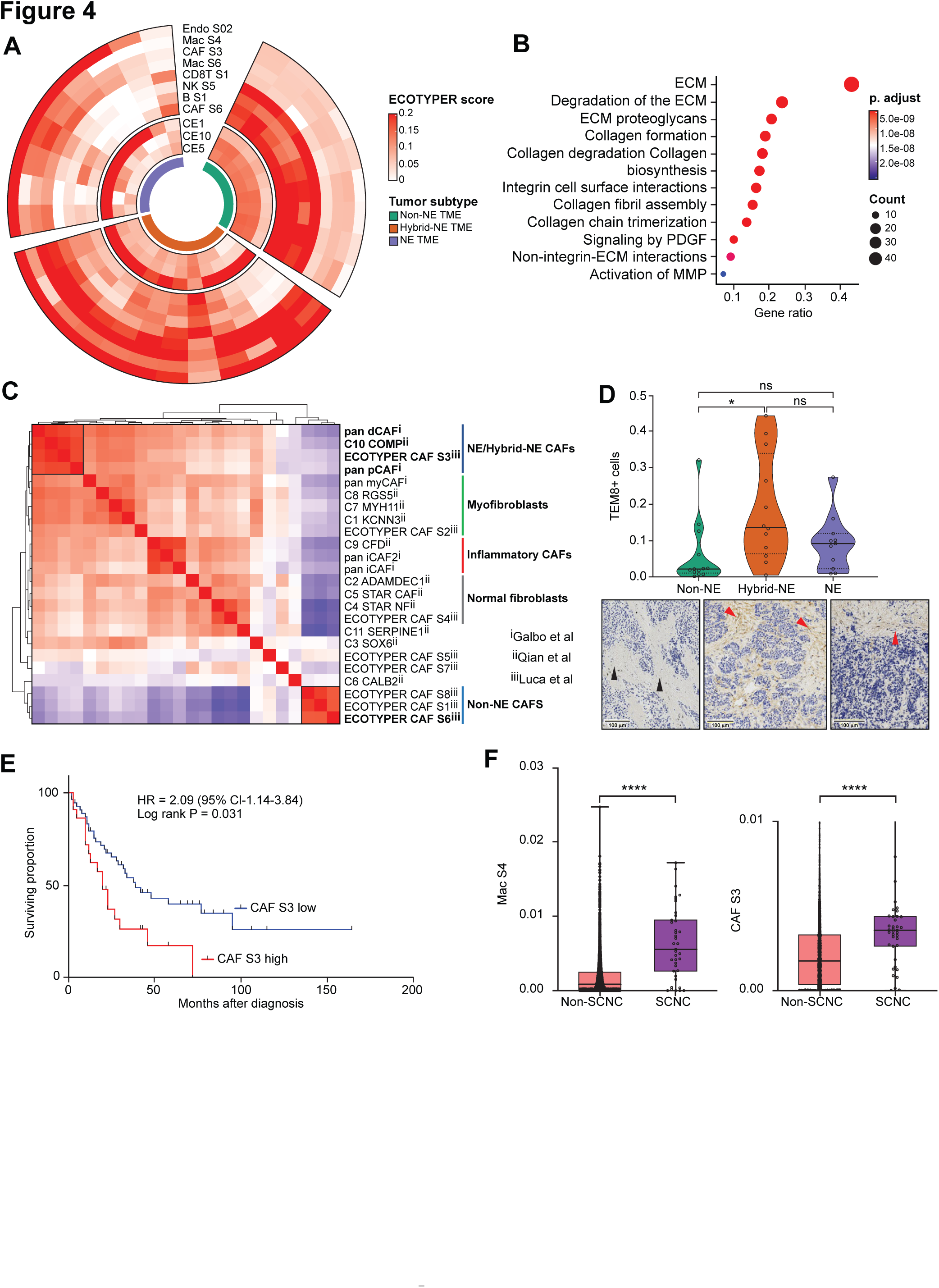
Immunosuppressive CAF cell state enriched in hybrid-NE SCLC. A. Differentially enriched cancer ecotypes and cell states across TME (n=30) subtypes^57^. B. Expression programs enriched in CAF S3. C. SCLC mesenchymal cells characterized using CAF signatures^57–59^ overlayed on sc-RNAseq data of mesenchymal cells extracted from the HTAN dataset^50^. Pair-wise correlation heatmap of ssGSEA-derived enrichment scores are shown. D. TEM8 protein expression by IHC (% TEM8 expressing cells) across TME subtypes (n=30) on sublevel sections of tumors profiled using spatially resolved transcriptomics^%^. Representative images (below) showing absence of TEM8 expression in non-NE TME (black arrows) and membranous and cytoplasmic expression in hybrid-NE and NE TME (red arrows). Color codes as for **Fig. 6A**. Scale bar set at 100µm. Also See figure S4E. E. Kaplan Meier curves showing survival (months) of patients with SCLC^5^ (n=81) with high and low CAF S3 expression (cut-off at 75% percentile). F. Enrichment of CAF S3 (right) and Mac S4 (left) in SCNC pan-cancer^38^.^#^ Abbreviations: SCLC, small cell lung cancer; TME, tumor microenvironment; ST, spatial transcriptomics; Endo, endothelial; CAF, cancer associated fibroblasts; Mac, Monocytes/macrophages; ssGSEA, single sample gene set enrichment; SCNC, small cell neuroendocrine carcinoma; OS, overall survival; HR, hazards ratio; CI, confidence interval; IHC, immunohistochemistry; TEM8, tumor endothelial marker 8; ECM, extra cellular matrix; MMP, matrix-metalloproteinases; PDGF-platelet derived growth factor; Diff, Difference; *statistical significance at p<0.05, **** statistical significance at p<0.0001; ^%^ Tukey’s multiple comparison test; ^#^ student t-test.

To contextualize the properties of CAF S3, we performed pairwise correlation of previously published CAF signatures, applying them to human SCLC mesenchymal cells^50^ (**Fig. 4C)** and spatially resolved TME segments of current study (Fig. S4C). This analysis revealed a striking similarity of CAF S3 with other aggressive CAF phenotypes such as pan-proliferative CAFs, pan-desmoplastic CAFs^58^ and C10 COMP CAFs^59^, which in previous studies have been associated with glycolysis, hypoxia, EMT, and metalloproteinase expression. While distinct, these CAF signatures shared expression of several genes (*TEM8, FN1, INHBA, POSTN* and *THY1*; Fig. S4D, Table S6) that have been individually associated with cancer stemness and tumor aggressiveness^60,61^. Tumor endothelial marker 8 (TEM8), also known as anthrax toxin receptor 1 (ANTXR1) is a highly conserved transmembrane receptor broadly overexpressed on CAF, endothelium, and pericytes. TEM8 is also a receptor for Seneca Valley virus, an oncolytic picornavirus, previously described to have selective tropism non-NE SCLC^62^. We profiled sublevel sections of the spatially profiled tumors using TEM8 IHC and confirmed the strong expression of TEM8+ cells in hybrid-NE TME (**Fig. 4D**). We also found heterogeneity of TEM8 expression in spatially separated areas within the same tumor, with expression limited to hybrid-NE TME (Fig. S4E; 2H,2I).

Importantly, SCLC patients^5^ whose tumors contained high proportions of CAF S3 had significantly worse outcomes [HR=2.09 (CI 95%-1.14-3.84, log-rank p=0.031)] compared to those with lower number of CAFs (**Fig. 4E**). CAF S3 remained an independent risk factor for death after controlling for clinical variables including age, gender, and stage (Fig. S4F). CAF S3 and Mac S4 (**Fig. 4F)** cell states were also enriched in pan-cancer small cell neuroendocrine tumors^38^, suggesting that these states may in part underlie the aggressiveness of small cell cancers regardless of tissue of origin.

Taken together, these analyses reveal CAF heterogeneity in SCLC TME, with hybrid NE subtype showing remarkable enrichment of CAF S3, marked by high expression of TEM8, with immunosuppressive and metastases aiding capabilities and portending poor prognosis.

### Proteomic characterization of tumor heterogeneity-linked reprogramming of SCLC TME

In parallel with spatially resolved transcriptomics, we used laser capture microdissection to separately enrich tumor and TME with the goal of proteomic characterization of SCLC heterogeneity (Fig. S5A). Fifteen tumor and 13 TME segments from rapid autopsy derived tumors of 11 patients were examined (see Methods, Table S7). Twelve tumors had matching bulk RNA-seq data, and six tumors had matching spatially resolved transcriptomic data described earlier(Fig. S5B). The histology-resolved tumor and TME were analyzed by quantitative MS-based proteomics^63^, resulting in 7418 and 6655 total proteins quantified in tumor and TME, respectively, and 6155 common proteins across both compartments (Fig. S5C). Enrichment of TME and tumor compartments was effective; tumor and TME samples separated widely on PCA for 1000 proteins with highest variance (**Fig. 5A**). TME segments were significantly enriched for stromal proteins (e.g., COL1A1, BGN, CD163) (**Fig. 5B**) and showed upregulation of stromal pathways (e.g., integrins, syndecan, and interleukin 4) (**Fig. 5C**). Tumor segments were significantly enriched for tumor-specific proteins (e.g., EPCAM, PARP1, EZH2), had higher proteomics-derived NE scores (Fig. S5D), and showed upregulation of tumor-related pathways e.g., DNA replication and repair (ATM, Fanconi, BARD1), tumor suppressors (RB1, E2F targets, P53 regulation) and MYC. As expected, tumor proteomes from different metastatic sites of the same patient were more like each other than tumors from other patients (Fig. S5E).

**Figure 5:**
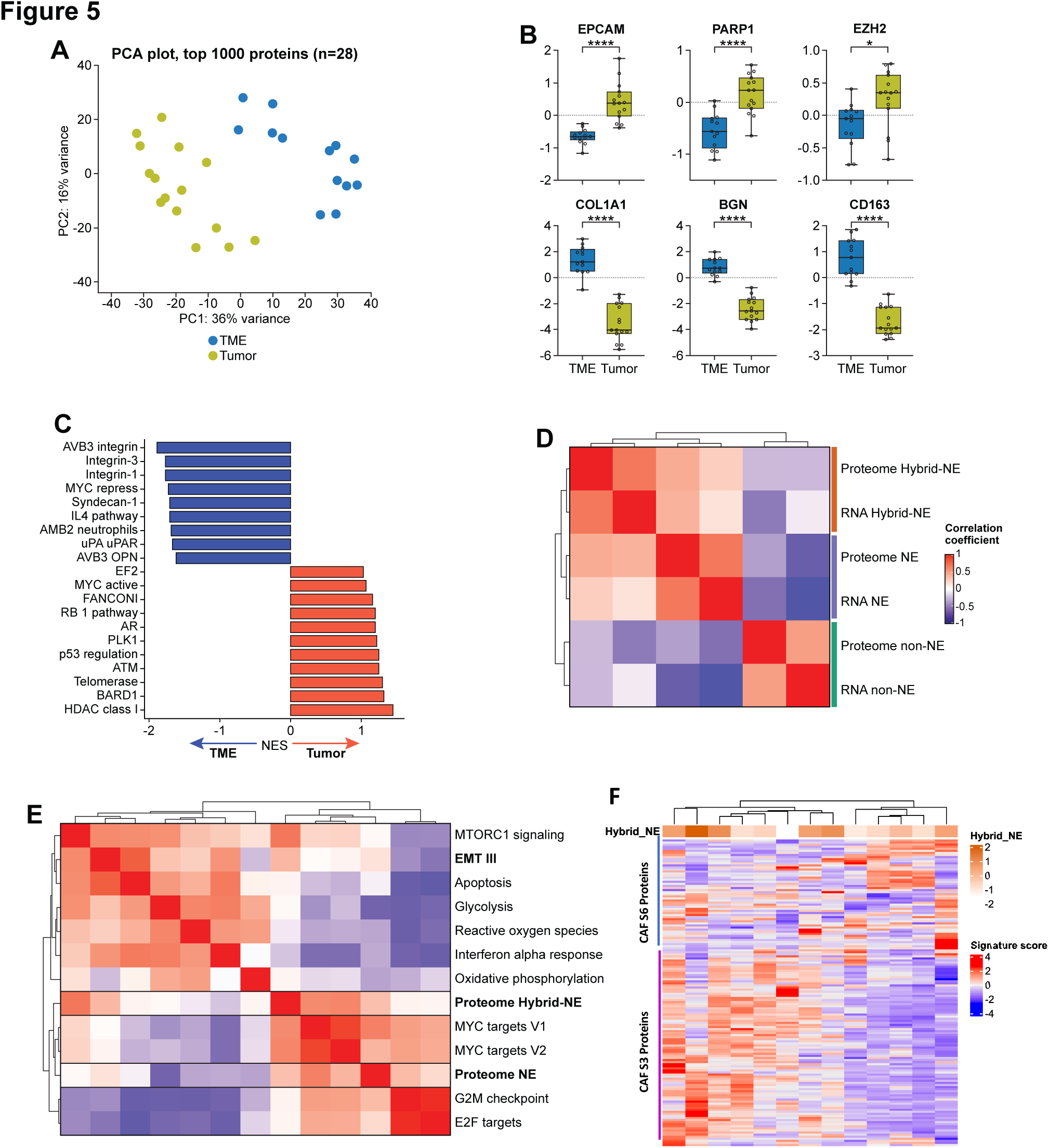
Proteomic characterization of tumor heterogeneity-linked reprogramming of SCLC TME. A. PCA of 1000 proteins with highest variance between tumor (n=15) and TME (n=13). B. Distribution of tumor (above) and TME (below)-associated proteins.^#^ C. NES (GSEA) of differentially expressed pathways between tumor and TME. D. Tumor transcript-protein correlation for NE and hybrid NE subtypes. Pairwise correlation of matched tumor proteome and bulk RNA-seq (n=12) derived NE, non-NE, and hybrid-NE signature scores. E. Pairwise correlation of tumor proteome-derived NE and hybrid-NE signatures with selected hallmark pathways (ssGSEA). F. Heatmap showing CAF S3 and CAF S6 proteins enrichment in TME proteome and their association with hybrid NE signatures of corresponding tumor proteome. Abbreviations: SCLC, small cell lung cancer; BGN, biglycan; EZH2, enhancer of zeste homolog 2; MS, mass spectrometry; PCA, principal component analysis; GSEA, gene set enrichment analysis; TME, tumor microenvironment; ssGSEA, single sample GSEA; NE, neuroendocrine; ns, non-significant at p<0.05; *statistical significance p<0.05; **** statistical significance at p<0.0001; ^#^ student t-test.

We then integrated proteomic data with patient-matched spatial transcriptomics data. A significant but modest correlation was observed between the overall tumor protein abundance and gene expression (Fig. S5F, median spearman correlation co-efficient 0.41; range, 0.11-0.47), in line with previously reported transcript-protein abundance correlations^64^. The correlation was more modest for TME segments (median 0.13; range, 0.05-0.26), possibly due to greater cellular heterogeneity of the TME compartment or reduced cellular density. In contrast to the modest correlation across all genes, we found strong correlation between proteomic and RNA-seq derived NE and hybrid-NE signatures (**Fig. 5D**, Fig. S5G). Proteome-based hallmark pathway analyses showed enrichment of DNA repair and replication stress pathways, E2F targets, and G2M checkpoints^10^ in NE SCLC. Hybrid-NE proteome was reminiscent of gene expression patterns described earlier (**Fig. 2H**), showing features of EMT (**Fig.5E**). Due to the low amount of available starting material and resultant low coverage of proteomes, only selected hallmark pathways with at least 50% coverage were included in these analyses (Table S8). TME proteome of hybrid-NE tumors showed enrichment of multiple CAF S3 compared with CAF S6 proteins (**Fig. 5F**). Additionally, TME proteome analyses supported earlier observations from spatially resolved transcriptomics including TAM-M2 enrichment and B/plasma cell de-enrichment with increasing NE differentiation (Fig. S5H) and CD8+ T cell exclusion in hybrid-NE enriched tumors (Fig. S5I). CD8+ T cell exclusion associated with hybrid-NE enrichment (Fig. S5J) was further confirmed in bulk RNA-seq datasets^9,16^.

Thus, histology-resolved proteomics confirmed key observations from spatially resolved transcriptomics, including the strong enrichment of hybrid-NE state with EMT-III, the aggressive CAF S3 subtype, and exclusion of CD8+ T cells. The high correlation between RNA and the corresponding protein levels suggests that RNA serves as a valuable indicator of protein expression for genes that contribute to SCLC NE heterogeneity.

### Tumor-TME crosstalk and tumor state modulation by FGFR inhibition

To determine spatially proximate communications that shape SCLC tumor cell states, we reconstructed cell-cell interactions based on coordinated expression of receptor-ligand pairs in TME and tumor^65^. We combined NE and hybrid-NE subtypes since TME of these two subtypes shared functional similarities (**Fig. 3B, 4A**).

Non-NE subtype exhibited strikingly higher number and diversity of interactions with TME compared with NE and hybrid-NE subtypes (**Fig. 6A**, S6A)^65,66^. Interactions significantly enriched in non-NE included immune checkpoint-receptor (e.g., PDCD1/PDCD1LG2), cytokine-receptor pairs (e.g., CCL15/CCR3)^9,43^, and multiple pathways of the FGF/FGFR signaling system especially FGF8 with multiple FGFR partners (**Fig. 6B**, Fig. S6B, Table S9). NE and hybrid-NE subtypes had far fewer interactions, but top interactions involved fibroblasts, macrophages, and endothelial cells further supporting the key role of these cell types in the NE and hybrid-NE ecosystems. These included MIF/CD74^67^, CD24/SIGLEC10^68^ and CD47/SIRPA^69^ signaling pathways. Unsupervised examination of differentially enriched programs between non-NE and NE/hybrid-NE TME corroborated with above observation, revealing multiple FGF/FGFR signaling pathways upregulated in non-NE TME (**Fig. 6C**, Table S10). To further validate our findings, we performed *FGF8* RNA in-situ hybridization on sublevel tumor sections (n=4). Highest *FGF8* signals were observed in non-NE TME (Fig. S6C). Consistently, we also found evidence of high FGFR activity in non-NE SCLC tumor segments supporting unusually high TME to tumor FGFR signaling (Fig.S6D).

**Figure 6:**
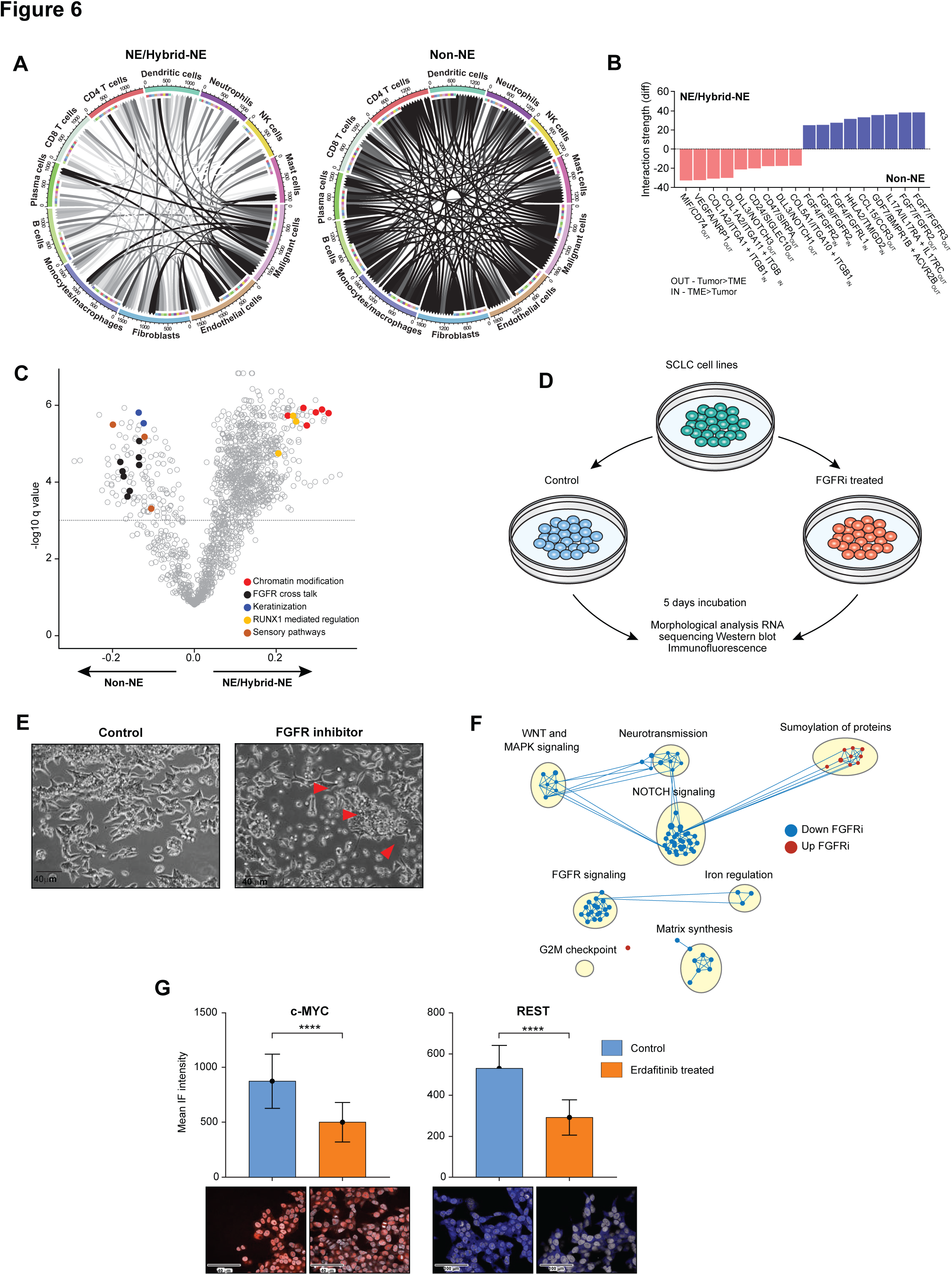
SCLC cell states modulated by FGFR inhibition. A. Intercellular tumor-TME interactions^57,65^ of NE/hybrid-NE (left) and non-NE tumor-TME (right) ecosystems. B. Differentially over-represented ligand-receptor interactions in NE/hybrid-NE and non-NE tumor ecosystems. Directionality of interaction specified for each pairs as in (TME-> tumor) and out (Tumor->TME). C. Differentially enriched expression programs (ssGSEA) between non-NE and NE/hybrid-NE TMEs. Recurrent and SCLC-relevant programs are highlighted. D. Workflow of FGFR inhibition in SCLC cell lines using pan-FGFR inhibitor erdafitinib. E. DMS-273 cell line treated with FGFR inhibitor (erdafitinib, 33.33 nM). Representative microscopic images (40x magnification) of untreated (left) and treated cells (right). Red arrows showing cells in suspension state. F. Pathways altered following treatment of DMS-273 with FGFR inhibitor (erdafitinib, 33.33 nM). Network enrichment plot of GSEA shown. Blue dots indicate pathways downregulated and red dots indicate upregulated pathways compared with control cells. G. c-MYC (left) and REST (right) mean IF intensity following treatment of DMS-273 with FGFR inhibitor (erdafitinib, 33.33 nM).^%^ Abbreviations: FGF8, fibroblast growth factor 8; RNA ISH, ribonucleic acid in situ hybridization; FGFR, fibroblast growth factor receptor ssGSEA; single sample gene set enrichment analysis, NE, neuroendocrine; nM/L, nanomoles/liter; IF, immunofluorescence; **p<0.01; ****p<0.0001; ns, not significant; R, Spearman’s correlation coefficient; ^#^Tukey’s multiple comparison test; ^%^ student t-test

Considering that FGFR signaling plays a pivotal role in the growth and differentiation of normal tissues and during embryogenesis^70^, we postulated that TME-derived FGF signaling might be crucial for maintaining the non-NE SCLC cell fate. It is important to note that while cell lines lack TME, cell culture media typically contain variable amounts of FGF in fetal bovine serum^71^. We treated non-NE SCLC cell lines NCI H211 and DMS-273 with varying concentrations of FDA approved pan-FGFR inhibitor erdafitinib for 5 days (**Fig. 6D**, Methods). Treatment with FGFR inhibitor led both cell lines to transition from predominantly adherent to suspension states (**Fig. 6E**), a growth pattern typically associated with NE differentiation^7,8^.

Network analysis of transcriptomic data revealed robust downregulation of various FGFR signaling pathways (**Fig.6F**), and well as non-NE signaling pathways such as MAPK^72^ and NOTCH signaling^29^, along with EMT. Correspondingly, western blot and immunofluorescence showed reduced expression of c-MYC which drives the non-NE cell fate^28^ and REST, a repressor of neural gene expression (**Fig. 6G**, Fig. S6E), indicating the dependence of SCLC non-NE cell state on extrinsic FGF signaling. Notably, there was no observable upregulation of NE markers at the protein level. This finding is reminiscent of irreversible fate switches induced by Notch^73^, a recurrent mechanism implicated in lineage switching in SCLC^29^. While Notch activation has been demonstrated to promote an NE to non-NE transition in SCLC, studies suggest that Notch blockade cannot completely reverse cells back to an NE-state^28,29^. Caspase activation and cell viability assays (Fig. S6F, S6G) did not reveal a significant increase in apoptosis with erdafitinib, indicating that the reduction in non-NE features is not attributable to apoptotic cell death.

## Discussion

SCLCs epitomize cancers with exceptional chemoresistance and metastatic capabilities, driven by significant intratumoral heterogeneity. Tumor-intrinsic factors that heighten SCLC plasticity include Notch pathway activation^29^ and MYC hyperactivation^28^ through amplification of extra chromosomal elements^30^. However, the cell-extrinsic determinants governing SCLC heterogeneity remain inadequately understood. Our comprehensive approach, integrating histopathology with spatially resolved transcriptomics and MS-based proteomics, whole-genome sequencing and multiplex immunofluorescence, maps SCLC tumor and TME states within their spatial context. Our work highlights the following key findings: (i) recognition of CAFs as a crucial element contributing to SCLC TME heterogeneity, with aggressive CAF subpopulations enriched in the hybrid-NE TME, predicting an exceptionally poor prognosis; (ii) substantial variation in the phenotype and overall composition of non-malignant cells in the metastatic TME, closely mirroring the tumor NE state, indicating TME-driven reprogramming of NE cell states. Notably, we find that TME-derived FGF signaling directs the tumor cells towards a non-NE cell state. Collectively, our work provides crucial insights into SCLC heterogeneity and the pivotal role of the TME in facilitating SCLC plasticity.

Previous studies have highlighted CAF heterogeneity in various solid tumors, characterized by differential marker expression and functional roles, including cancer invasion and metastasis through secretion of soluble factors and matrix remodeling^74^. CAFs have not yet been studied in detail in metastatic and relapsed SCLC. largely due to limited tissue availability. In the present study, profiling of rapid autopsy-derived tumors enabled the discovery of CAF as a major component of the SCLC TME. CAF subpopulations with heterogenous marker expression were identified even within the same tumor. The complexity and heterogeneity of CAFs appear to be closely interconnected with the tumor neuroendocrine state. Particularly, the hybrid-NE TME is characterized by an abundance of FAP/POSTN-expressing CAFs, akin to previously described CAF subtype S3^57^, the oncofetal CAFs with immunomodulatory properties^75^ and recently described immunosuppressive senescent CAFs^76^. These CAFs exhibit enrichment for extracellular matrix, collagen organization and degradation pathways. Correspondingly, we observed pronounced stromal expression of both nuclear and cytoplasmic YAP1 in hybrid-NE SCLC, consistent with the recognized role of YAP1 in the establishment and maintenance of CAFs^77^, as well as its activation in cancer cells in close proximity to a rigid matrix^77,78^. The factors driving CAF heterogeneity and the implications of CAF infiltration in SCLCs, particularly regarding their immune regulatory functions, warrant further investigation. Our findings align with a recent study that reported that SCLC display the highest FAP expression compared to various other solid tumors^79^.

Targeting the distinctive immunosuppressive mechanisms inherent to each cell state holds promise for reinstating immunosurveillance and enhancing SCLC responses to immunotherapy. This approach is particularly pertinent in SCLC, which, despite exhibiting a highly mutated genome, displays only modest responses to immunotherapy^80,81^. TME-targeted therapies may be broadly relevant for neuroendocrine cancers across tissues of origin, which are enriched for immunosuppressive CAFs. While early efforts to target CAF-cancer cell interactions in certain cancer types have hinted at the potential drawbacks of stromal ablation, such as the development of more aggressive cancers^82,83^, a nuanced understanding of CAF diversity and their influence on immunosurveillance could inform the development of personalized treatment strategies for SCLC.

Our work provides crucial insights into the role of the TME in shaping the SCLC phenotype, an area that has remained poorly understood primarily due to the limited availability of patient tumor samples for research. Tumor samples obtained through rapid research autopsies from patients with metastatic SCLC provide the most compelling evidence to date of extensive transcriptional heterogeneity heavily influenced by the TME. In contrast to the NE and hybrid-NE subtypes, we find intimate interactions between the non-NE subtype and the adjacent TME, actively reprogramming the tumor states. Notably, previous studies in model systems have implicated FGF signaling as a crucial component mediating the cross-talk between NE and non-NE cell states^8,84^, as well as a defining feature of chemotherapy-resistant persister cells^85^. Our findings demonstrate that, in human tumors, FGF signaling can originate from the TME, sustaining a chemo-resistant non-NE state. These results align with the previously established role of FGF signaling in driving SCLC development, particularly non-NE cells^86^, and offer an explanation for the observed sensitivity of chemo-resistant non-NE SCLC cells to FGFR inhibition^87,88^. Furthermore, a dependency on FGF signaling has been reported in a subset of castration-resistant prostate cancers characterized by limited or absent androgen receptor expression and non-NE features^89^. Collectively, these findings underscore TME-derived FGF signaling as a shared pathway underlying the lineage plasticity of NE cancers across various contexts and may guide future endeavors to modulate tumor-TME crosstalk and restrain tumor evolution.

### Limitations of the study

This study represents a comprehensive report of tumor-extrinsic drivers of SCLC plasticity in human tumors and provides insights into CAFs as key TME elements in SCLC. Nevertheless, several notable limitations should be acknowledged. These include the lack of single-cell resolution at a spatial level inherent to the GeoMx NanoString approach, which prevents definitive conclusions regarding the precise localization and composition of the hybrid NE state. We also recognize that this manuscript has not extensively validated the functional impact of the observed hybrid-NE state and the surrounding CAFs, including their impacts on chemo-resistance and metastasis. The study is limited by the relatively small number of samples, but the validation using orthogonal proteomic based do strengthen our conclusions. Moreover, our dataset represents heavily treated, relapsed SCLC tumors and so the generalizability of these findings in untreated early-stage SCLC and treatment naïve extensive stage SCLC needs to be confirmed. Additionally, further studies using tumors from a wider range of metastatic sites are needed to understand how the TME programs are influenced by the site of metastasis.

## Acknowledgement

We gratefully acknowledge the contributions of our patients and their families. This work utilized the computational resources of the NIH HPC Biowulf cluster (http://hpc.nih.gov). We thank NanoString GeoMx’s early access whole transcriptome atlas program (Jingjing Gong, Stephan Phelan).

## Author Contributions

Conceptualization, P.D., N.T., T.C., A.T.; Methodology, P.D., R.K., J.M., A.H., C.S., Y.C., D.T., D.N., L.C., K.J., V.R., M.T., D.L., Y.Z., P.F., N.W.B., T.A., G.N.; Investigation, P.D., S.N., A.T., L.P., A.S., S.K., B.S., A.K.S., T.C., R.V., D.P., C.G., D.B., A.W., R.S., M.M., K.B., J.C., A.B., R.S., M.A., M.K., G.C., M.N., S.G., D.A., G.T.B., M.K.J., D.K.,A.S., S.H.;R.E.M.; Supervision, L.S.,A.A., U.G.,Z.O.W.,E.R.,M.M.,A.T.; Writing-Original Draft, P.D., N.T., A.T.; Writing-Review & Editing, P.D., A.T.; Resources-S.H.,A.T.

## STAR Methods

### Lead Contact

Further information and requests for resources and reagents should be directed to and will be fulfilled by the lead contact, Anish Thomas.

### Materials availability

This study did not generate new unique reagents.

### Resource availability

Processed spatial transcriptomics data (**Extended data 1**), whole genome sequencing mutation and copy number call out (**Extended data 2**), mass spectrometry proteomic data (**Extended data 3**), Bulk RNA-seq data of tumors profiled for proteomics (**Extended data 4**) and DMS273 cell line data using FGFR inhibitor (erdafitinib at 33.33nM concentration) and control (untreated cell line), (**Extended data 5**) are available with this manuscript. Additionally, above raw and processed spatial transcriptomics and RNA-sequencing data is also submitted on public database on GEO as GSE267310. The mass spectrometry proteomics data have been deposited to the ProteomeXchange Consortium via the PRIDE partner^91^ repository with the dataset identifier PXD052033. This paper does not report any original code. Any additional information including microscopy data required to reanalyze the data reported in this paper is available from the lead contact upon request.

## Experimental Models and Study Participant Details-

### Human Samples-

***NCT01851395*** is approved by NIH Institutional ethics committee and Institutional review board (IRB). Detailed informed consents were obtained from all patients and their next of kin (after demise of the subject) as per protocol *NCT01851395.* Rapid autopsy was performed at NIH Clinical center laboratory of Pathology (LP) Autopsy suite by LP pathologists. Tissue was collected, stored and processed as per the detailed plan laid out in protocol NCT01851395.

Samples for spatial transcriptomics and TME profiling were obtained from the first 10 consecutive patients who died of metastatic small cell lung cancer and were enrolled into the rapid autopsy protocol ***NCT01851395.*** Multiple sampled tumor sites from every patient were studied for the presence or absence of necrosis and/or autolytic changes and tumors with the least amount of these and preserved morphology were selected for spatial profiling. One tumor was selected per patient. Clinical details were recorded (table S1). Multiple 4–5-micron serial sections from each tumor FFPE block were taken. The first two sections were processed for spatial transcriptomics profiling (see below), immediately followed by separate unstained sections for mIF, IHC and mass spectrometry-based proteomics in that order. Following these sections, DNA extraction (below) for whole genome sequencing as well as bulk RNA sequencing was performed wherever feasible.

15 samples were obtained from 11 unique patients who died of metastatic small cell lung cancer and were enrolled into the rapid autopsy protocol ***NCT01851395*** for mass spectrometry-based proteomics profiling. Table S7 includes important clinical and site information regarding these samples. Three patients had >1 tumor sample profiled, and 6/15 tumor samples profiled had matched spatial transcriptomics data as profiled above albeit on a different serial section. Majority (12/15) of the tumors had matched bulk RNA-seq profiled and were used to find gene-protein correlation.

### Cell-lines

Briefly, established SCLC cell lines (DMS 273 and NCI H211) were grown in RPMI-1640 media supplemented with 10% FBS, Pen-Strep and L-Glutamine at 37°C and 5% CO2. Cell aliquots were treated with either DMSO or FGFR inhibitor (Erdafitinib) dissolved in DMSO dosed at 3.3, 10 and

33.33uM concentrations once at the beginning of a 5-day interval period. At the end of 5 days, cells were subsequently analyzed for morphological analysis, RNA extraction using standard pipelines for RNA sequencing, and cell lysates preparation for western blot staining.

## Methods Details-

### Spatial transcriptomics experiment and analysis

GeoMx^R^ Human Whole Transcriptome atlas (GeoMx Hu WTA) was used for the tumors selected for spatial transcriptomic analysis under a special early access program. Two consecutive sections of 4µm – thick slides were prepared from parent FFPE blocks. 1^st^ slide was H&E stained to visualize the tumor and TME regions. Random but non-necrotic and non-hemorrhagic regions of interest (ROI) were selected by a board-certified pathologist. 2^nd^ slide was deparaffinized, heated in ER2 solution (Leica) at 100 degrees C for 20 minutes, and treated with 1 microgram/ml of proteinase K (Ambion) at 37 degrees C for 15 minutes on a BOND Rxm autostainer (Leica). An overnight in situ hybridization was performed with a probe concentration of 4nM per probe as described previously^92^. Slides were washed twice at 37 degrees for 25 min with 50% formamide/2X SSC buffer to remove unbound probes. Prepared slides were stained further with pan-cytokeratin (AE1+AE3, Novus Biologicals,1:500) for epithelial/tumor cells, CD45 for immune cells (D9M8I, CST, 1:100), Syto83 for nucleus (stainSyto83, Thermofisher, 1:25). Stained slides were loaded onto GeoMx instrument and scanned. Forty (40) circular ROIs of 500µm diameter were selected randomly across different areas of 10 tumor slides (**see Extended Data 6**). Using the information from pan-CK and morphological features, tumor cells/areas were selected and marked as “pan-CK positive/Tumor” areas, and the rest of the area was marked as “pan-CK negative/TME” areas to perform segmentation of most of the ROIs(n=32/40) where the clear distinction of “TME” and “Tumor” segments was possible. In addition, 4 regions with no/minimal visible TME were profiled as “tumor only” segments and other 4 regions without any tumor in near distance were profiled as “normal” segments.

After sequencing, reads were trimmed, merged, and aligned to retrieve the probe identity. The unique molecular identifier region of each read was used to remove PCR duplicates and duplicate reads, thus converting reads into digital counts. The sequencing saturation was sufficient for RNA and was >80% for all the ROIs. For each gene in each sample, the reported count value is the mean of the individual probe counts after the removal of outlier probes. The limit of quantification (LOQ) was set at the geometric mean plus two standard deviations of the negative probes. 18676 genes were targeted and captured by DSP. Logarithmic (base 2) conversions of normalized third Quantile expression values were used for downstream analysis. The Euclidean distance metric was used for principal component analysis (PCA).

### Whole genome sequencing (WGS) and Bulk RNA sequencing

DNA and RNA were extracted from FFPE tumor tissues used for spatial and proteomics profiling for WGS and bulk RNA-sequencing using standardized DNA extraction kits. Matched normal DNA was extracted from stored blood. Similarly for cell lines RNA extraction was performed for cell lines (DMS-273) exposed to DMSO and at 33.33uM FGFRi concentration (4 technical replicates each). Samples were pooled and sequenced on Novaseq 6000 using S4 flow cell configuration using Truseq Nano DNA library prep and 150bp paired end sequencing was done. All the samples have percent of Q30 bases above 88%. All the samples have yields between 193 and 389 million pass filter reads. Human DNA libraries were sequenced with the aim of obtaining coverage of a minimum 70X for tumor DNA and 30xmatched normal DNA. QC and Alignment –For all whole genome and whole exome data, raw FASTQ reads were trimmed for adapter and low-quality control using Trimmomatic1 (v. 0.33) prior to alignment^93^. Alignment was performed using BWA2 (v. 0.7.17) mapping to the human reference hg38 genome. Duplicated reads were marked using Picard (v. 2.17.11) followed by indel realignment and base quality score recalibration using GATK (v. 4.2.2.0)^94^. Variant Calling and Annotation-Somatic variant calling was performed using Mutect2, in both tumor-normal and tumor-only mode using the GATK best practices (v. 4.2.2.0). Variants were annotated using the Ensembl Variant Effect Predictor and the vcf2maf tool (v. 102)^95^. MAF files were used for all downstream annotation and visualization. Variants were filtered removing variants with tumor read depths <5, an alternate allele count <2, and normal alternate allele counts >1. Additionally, to filter commons polymorphisms, variants were removed with a frequency >0.001 in the ExAC, gnomAD, or 1000 Genomes population databases. Copy Number Variant Analysis-For all samples, cn.MOPS4 was used for CNV calling. Aligned and processed BAM files were converted to read count matrices used as input. The software models the depths of coverage across samples at each genomic position to account for read count biases along chromosomes. For WES samples, the exomecn.mops function to adjust for the varying window spans across the target regions. RNA was extracted from FFPE tumor cores (n= 12 samples) using RNeasy FFPE kits according to the manufacturer’s protocol (QIAGEN, Germantown, MD). RNA-seq libraries were generated using Truseq RNA Access Library Prep Kits (TruSeq RNA Exome kits; Illumina) and sequenced on Highseq3000 sequencers using 75□bp paired-end sequencing method (Illumina, San Diego, CA). For transcriptomic analyses, raw RNA-Seq count data were normalized for inter-gene/sample comparison using FPKM, followed by log2(x□+□1) transformation, as implemented in the edgeR R/Bioconductor package^96^

### Multiplex Immunofluorescence (mIF)-

mIF was performed through our collaboration with the Human Immune Monitoring Shared Resource (HIMSR) at the University of Colorado School of Medicine. We performed 6 colors multispectral imaging using the Akoya Biosciences Vectra Polaris instrument. Unstained FFPE-derived slides were on the Leica Bond RX autostainer according to standard protocols provided by Leica and Akoya Biosciences and performed routinely by the HIMSR. Slides from spatially profiled tumors were stained consecutively with antibodies specific for the following proteins: antibodies (see above table) and DAPI counterstain. Briefly, the slides were deparaffinized, heat treated in antigen retrieval buffer, blocked, and incubated with primary antibody, followed by horseradish peroxidase (HRP)-conjugated secondary antibody polymer, and HRP-reactive OPAL fluorescent reagents that use TSA chemistry to deposit dyes on the tissue immediately surrounding each HRP molecule. To prevent further deposition of fluorescent dyes in subsequent staining steps, the slides were stripped in between each stain with heat treatment in antigen retrieval buffer. Whole slide motif scans were collected using the 20x objective. The 6 color images were analyzed with inForm software to unmix adjacent fluorochromes, subtract autofluorescence, segment the tissue, compare the frequency and location of cells in the tumor and stromal areas, to segment cellular membrane, cytoplasm, and nuclear regions, score each cellular compartment for expression of any scoring markers you include in your panel, and phenotype infiltrating immune cells according to morphology and cell marker expression. Segments that were profiled using GeoMx spatial transcriptomics and retained the tumor and TME morphology on the sublevel section (as deemed by board certified pathologist) used for mIF were marked and quantified for cells/density parameter.

### Immunohistochemistry-

All stains were run on our Leica Bond Max auto-stainer with standard DAB protocol. IHC stains (antibodies listed in the resource table) were performed at the National Institutes of Health (NIH), Laboratory of Pathology and at Molecular histopathology laboratory, Frederick national laboratory, NCI according to the manufacturer’s instruction. IHC-stained slides were scanned using the Carl Zeiss AxioScan.Z1 microscope equipped with a Plan-Apochromat 20x NA 0.8 objective. For Nuclear markers-H-score was calculated based on the equation: 1 x (% of weakly stained nuclei) + 2 x (% of moderately stained nuclei) + 3 x (% of strongly stained nuclei). For Cytoplasmic and Membranous markers % positive cells out of total cells in a given segment (based on nuclei scoring) were used to report. Segments that were profiled using GeoMx spatial transcriptomics and largely retained the tumor and TME morphology on the section (as deemed by board certified pathologist) used for IHC were marked and quantified for cells/density parameter. Nuclear and cytoplasmic IHC scoring was done using Qu-Path (open source v0.3.0) software.

### Mass-spectrometry based quantitative proteomics-

Consecutive tissue thin sections (8 μm) were generated by microtome, placed onto polyethylene naphthalate (PEN) membrane slides, and stained using hematoxylin and eosin (H&E). Cellular enrichment via laser microdissection (LMD) was performed on the LMD7 (Leica Microsystems) for selective harvest of tumor and TME regions from each of the 15 tissue specimens (**See extended data 7**). LMD enriched samples in 20 µl 100 mM triethylammonium bicarbonate / 10% acetonitrile were pressure-assisted, trypsin-digested using pressure cycling technology (PCT; Pressure Biosciences, Inc.), as previously described^63^. Briefly, LMD samples were incubated at 99 °C for 30 min followed by 50 °C for 10 min with SMART trypsin (ThermoFisher Scientific). SMART trypsin was added at a ratio of 1 μg per 30 mm^2^ tissue. Lysis and digestion were performed in a Barocycler 2320EXT (Pressure Biosciences, Inc.) by cycling between 45 kpsi for 50 sec and atmospheric pressure for 10 sec, for 60 cycles at 50 °C. Peptide digests were quantified by colorimetric assay (Pierce BCA Protein Assay Kit). Digested peptides (10 µg) from each sample (n=15 LMD enriched tumor and n=13 LMD enriched TME samples) were labeled using isobaric tandem mass tags (TMT), per manufacturer’s instructions (TMTpro 18-plex Isobaric Label Reagent Set, ThermoFisher Scientific)^97^. A reference sample was generated by pooling equivalent amounts of peptide digests from each of the patient samples and included in each TMTpro-18 multiplex. TMT-labeled samples were pooled within respective multiplexes, cleaned using the EasyPep Maxi MS Sample Prep Kit (ThermoFisher Scientific) and fractionated offline to 36 pooled fractions via basic reversed-phase liquid chromatography (bRPLC). The TMTpro-18 bRPLC fractions were analyzed by liquid chromatography tandem mass spectrometry (LC-MS/MS) employing a nanoflow LC system (EASY-nLC 1200, ThermoFisher Scientific) coupled online with a Q Exactive HF-X MS (ThermoFisher Scientific), as previously described^63^.

### RNA Insitu-Hybridization

FGF8 expression was detected by staining 5 um FFPE tissue sections with RNAscope® 2.5 LS Probe Hs-FGF8-C2 (ACD, Cat# 415798-C2) and the RNAscope LS Multiplex Fluorescent Assay (ACD, Cat# 322800) using the Bond RX auto-stainer (Leica Biosystems) with a tissue pretreatment of 15 minutes at 95°C with Bond Epitope Retrieval Solution 2 (Leica Biosystems), 15 minutes of Protease III (ACD) at 40°C, and 1:750 dilution of TSA-Cyanine 3 Plus (AKOYA). The RNAscope® 3-plex LS Multiplex Negative Control Probe (Bacillus subtilis dihydrodipicolinate reductase (*dapB*) gene in channels C1, C2, and C3, Cat# 320878) was used as a negative control. The RNAscope® LS 2.5 3-plex Positive Control Probe-Hs was used as a technical control to ensure the RNA quality of tissue sections was suitable for staining. Slides were digitally imaged using an Aperio ScanScope FL Scanner (Leica Biosystems). All image analysis was performed using HALO imaging analysis software (v3.4.2986.246; Indica Labs, Corrales, NM), and image annotations were performed by one pathologist (BK). The analysis was performed using FISH V3.1.3 in HALO to determine percent positive FGF8 positive cells. Areas of artifact such as folds, and tears were excluded from analysis.

### Western blotting

Cell lines (DMS 273 and NCI-H211) were exposed to FGFR inhibitor at 0 (control), 3.3, 10 and 33.33nM concentrations for 5 days and washed with PBS and lysed in lysis Buffer and protease inhibitors. Protein (30-50 microgram) was loaded into gels and overnight transfers were performed and blots were probed with antibodies against MYC, Erk1/2, pERK ½, REST, INSM1, NEUROD1, YAP1 and with ACTIN (Millipore Sigma) as a loading control (see resource table for more information). These procedures were repeated twice.

### Cell viability assay

DMS 273 and NCI-H211 cells were plated at 1,000 cells per well in 384 well plates and treated with erdafitinib at the concentrations indicated above. After 5 days cells were collected per manufacturer’s instructions using either CellTiterGlo® (Promega, G7570) to measure viability by assessing ATP concentrations, or using Caspase-Glo® 8 Assay Systems® (Promega, G8200) to measure cleaved Caspase 8 activity.

## Quantification and Statistical Analysis

R studio version 4.1.2 (2021-11-01) (R Foundation for Statistical Computing), GraphPad Prism version 9.0.2 (GraphPad Software) and Qlucore Omics Explorer version 3.7 (Qlucore AB) were used to generate figures and perform statistical analyses. Partekflow version 10.0.22.0321 (Partek®) was used for processed single cell sequencing analysis and visualization. All tests were two-tailed and p-values less than 0.05 were considered significant. False discovery rate (FDR) of 5% was used wherever applicable. Student’s t-test was used to compare between two groups and one-way Anova test followed by Tukey’s multiple comparison test was performed to compare numerical data between more than 2 groups unless otherwise specified.

### Spatial transcriptomics analysis-

Log(10) normalized data was used for subsequent analysis. Two TME segments that did not meet the sequencing quality metrics (Fig. S1C, segment 1007 and 1008) were excluded from further analyses.

### Optimal Cluster determination in spatial transcriptomics dataset (tumors)-

In order to determine ideal number of clusters amongst the 36 tumor segments profiled we calculated “total within-cluster sum of squares” values for projected cluster numbers (k) from 1-10 using the *nbClust* R package^98^. Based on this, we found that partitioning the data into 3 (k=3) clusters would be the best approach in order to avoid over-clustering, as determined by the “elbow” of the curve. Subsequently using k=3, we performed k-means unsupervised partitioning clustering^99^ (Iterations=50, Attempts=200) in order to annotate each tumor segment to a cluster.

### Cluster-defining genes determination-

For the tumor segments (n=36), 6000 highly variable genes were extracted Given that 3 well-defined clusters were visualized, PC1 and PC2 contribution in terms of coordinates axis of each gene was computed. positive contributors to PC1 were classified as cluster 1-specific signature (351 genes), the negative contributors to PC2 as the cluster 3-specific signature (149 genes), and the positive contributors to PC2 as the cluster 2-specific signature (244 genes).

### Quantitative proteomics data processing pipeline-

Peptide identifications, normalization, and log_2_-transformation for generation of protein-level quantitative data were performed, as previously described^63^. Briefly, raw data files were searched using Mascot (Matrix Science) and Proteome Discoverer (ThermoFisher Scientific, Inc). with a publicly available non-redundant human proteome database (Swiss-Prot, Homo sapiens, Proteome UP000005640, 20,257 sequences, downloaded 12-01-2017; http://www.uniprot.org/) appended with porcine trypsin (Uniprot: P00761) sequences. Peptide spectral match (PSM) results were filtered using a <1.0% false discovery rate (FDR). Log_2_-transformed PSM abundances were calculated using z-score transformation for each TMT-18 channel relative to the pooled reference standard. Protein-level abundances were calculated from the normalized log_2_-transformed TMT reporter ion ratio abundances from ≥2 PSMs corresponding to a single protein accession. Protein-level abundances were imputed for proteins absent from individual patient samples, but present in ≥50% of all patient samples.

### Survival analysis-

Ecotyper cell state scores were computed for bulk RNA sequencing data (George et al). For Ecotyper_CAF_S03 scores, an arbitrary cut-off of 75% percentile was used to divide samples into “low CAF_S03 (<75 percentile)” and “high CAF_S03 (>75 percentile). Survival analysis was further done using the cox proportional hazards model and log-rank testing was done to calculate the survival distributions of the two types of samples. Statistical significance was noted at value <p. Multivariate analysis was done after factoring in other available and potentially confounding variables (age, gender, smoking status).

### Tumor budding/nesting assignment-

All the tumor segments profiled (n=36) were analyzed for cyto morphological features on the initial Hematoxylin and Eosin stain. Tumor nesting was identified as feature where multiple tumor cell clusters were found to be surrounded by dense stromal tissue as previously reported^47^. Tumor budding was identified when tumor cells either single cells or small clusters (<5-10 tumor cells) were identified in a segment.

### Pairwise correlation analysis-

Normalized cell line and human bulk RNA data were downloaded and subsequently, ssGSEA based scores were computed for relevant gene signatures for each sample. Correlation metric pairwise correlation was done either for ssGSEA-derived gene sets or single normalized genes. Subsequently, a heatmap plot was constructed using Pearson’s correlation coefficients, which was then hierarchically clustered to observe the correlation patterns between genes/gene sets of interest.

### Tumor purity estimates-

For prior datasets^5,9^ where matched bulk RNA-seq and Whole genome/exome sequencing data was available tumor purity scores were estimated using ABSOLUTE^41^ algorithm. Subsequently ssGSEA derived stromal and immune scores^40^ were computed for these transcriptomes. Using these two scores a linear model was generated. Subsequently, stromal and immune scores were calculated for tumor segments of spatial transcriptomics profiled segments and tumor purity estimates were predicted using the above model.

### Single sample gene set enrichment analysis (ssGSEA)-

ssGSEA enrichment scores were computed using GSVA R/Bioconductor package^100^. Additionally, data generated for Figure 2C was calculated using ssGSEA scores for each tumor segment for the 50 hallmark gene sets downloaded from MsigDB. Subsequently, for each hallmark set, q value was calculated by comparing one cluster segments vs other two segments using two-way Anova followed by controlling for FDR method of Benjamini and Hochberg.

### Gene set enrichment analysis (GSEA)-

For each cluster, the marker genes were identified by comparing the cluster with the other clusters using the Wilcoxon rank-sum test. Then, the fold change in expression of each gene was computed by taking the taking the log2(median Q3 expression / median Q3 expression in other clusters). Genes lists were constructed using those genes with p-values□<□0.05 and an absolute fold change of >2. Enriched hallmarks from the Molecular Signatures Database were identified by pre-ranked GSEA using clusterProfiler v.4.0.5 using the gene list ranked by log-transformed p-values with signs set to positive/negative for a fold change of >1 or <1, respectively.

### Transcription factors downstream target activity-

An SCLC context-specific gene regulatory network was computed using the ARACNe-AP software^101^. Its associated transcription factor (TF) target sets (regulons) were then input to the VIPER software package^102^, to infer a TF x sample matrix of activity values. Each TF activity value is a normalized measure of regulon enrichment among sample-specific over/under expressed genes. ARACNe-AP was run using a bulk RNA-Seq data set derived from 127 SCLC tumor samples^9^ excluding genes with zero expression across all samples. The regulatory network was restricted to TFs and TF co-regulators associated with the Gene Ontology terms GO:0003700 (DNA-binding transcription factor activity) and GO:0003712 (transcription coregulator activity), together with their inferred targets (using the ARACNe-AP ‘--tfs’ parameter to specify the starting gene regulator set). The final network was derived by integrating 100 intermediate networks, constructed using bootstraps of the expression data set, as described^101^. The p-value for mutual information significance, a parameter governing the number of associations in the network, was set to 10^-8.The VIPER activity matrix was computed using the viper R/Bioconductor package (https://www.bioconductor.org/packages/release/bioc/html/viper.html). In particular, the viper::viper::aracne2regulon () function was used to derive a regulon object from the ARACNe-AP output and gene expression data, using the parameter setting “format=’3col’”. Activity data were then computed using latter regulon and the expression data as input to the viper::viper () function, with parameter settings ‘pleiotropy = TRUE’ and ‘nes = TRUE’.

### TME cluster assignment-

Each of the TME segment was assigned the cluster class of the tumor segment from which it was segmented (spatially proximate TME). Because TME segments of 2 regions (1007 and 1008) yielded low quality data, they were excluded from further analyses. Total of 30 TME segments were thus analyzed downstream.

### CIBERSORT deconvolution-

CIBERSORT tool developed by Newman et al^53^ to quantify cell types in bulk derived RNA expression data was used to predict cell types in TME segments. The analysis was run on the CIBERSORT website at http://cibersort.stanford.edu. We applied safeTME^103^ to the TME segments gene expression data. For each run, 100 permutations were performed, and quantile normalization was disabled.

### Cellular ecotypes and cell states computation and evaluation-

ECOTYPER is a recent tool developed by Luca et al^57^ to deconvolve tumor-microenvironment cell types and interactions between them as “multi-cellular ecotypes”. We ran the analysis on ECOTYPER website https://ecotyper.stanford.edu/carcinoma/ for TME segments. Both ecotypes and different cell states for each cell-type information was downloaded from the analyzed results and for each given TME segment a given Ecotype and a cell state with maximum enrichment was binned to that category. For the pan cancer tumors analysis, Ecotyper data for TCGA cell state assignments were downloaded from https://ecotyper.stanford.edu/carcinoma/ website. SCN scores of the TCGA tumors were obtained from publicly available data available with Balanis et al^38^.

### Cell-cell communication /ligand-receptor interaction-

We used ICELLNET^65^ to score sender-receiver (either TME segment->Tumor segment or Tumor segment->TME segment) expression profiles using a custom set of ligand-receptor (L/R) annotations from ECOTYPER^57^. Next, we computed a summary communication score for each sender-receiver cell population (as available in ECOTYPER) by taking the maximum ICELLNET score of all L/R pairs for each sender-receiver cell population. To confirm, findings from ICELLNET we also computed similar scores using an orthogonal pipeline CellphoneDB^66^. Briefly, Cellphone was run with the “cpdb_statistical_analysis_method”. Ligand-receptor interaction means, and p-values were generated, and ligand-receptor pairs were cross matched with annotations from ECOTYPER^57^ converting gene-gene interaction to cell-cell interaction. Lowest ligand/receptor interaction values were assigned to each cell/cell interaction after which a mean score was computed and plotted as a heatmap.

### Shannon diversity scores-

Shannon biodiversity index measures the diversity of a population distribution. Higher biodiversity could indicate the presence of a higher number of species, or homogeneity in the distribution of a smaller number of species. We define the Shannon index as:

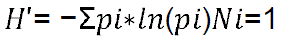

Here, N is the total number of species and *pi* is the proportion of i’th species.

### Additional Resources

**Rapid autopsy protocol link**: URL https://clinicaltrials.gov/ct2/show/NCT01851395

**Natural history protocol link**: URL https://clinicaltrials.gov/ct2/show/NCT02146170

## Supplemental Files

Supplemental table S1: Clinical and Pathological characteristics of spatially profiled tumors related to Figure 1A.

Supplemental table S2: Immunohistochemistry derived H scores for each segments of spatially profiled tumors related to Figure 2G.

Supplemental table S3: Transcript derived pathway scores for tumor segments related to Figure 2H.

Supplemental table S4: Cluster specific genes derived from PCA distribution for each of the 3-tumor cluster segments related to Figure 2I.

Supplemental table S5: Ecotyper scores for cellular ecotypes (CE) and various ecotype constituent cells related to Figure 4A.

Table S6: Cancer associated fibroblasts enriched in Hybrid-NE/NE segments associated gene signatures from various previously published groups related to Figure 4C and Figure S4D

Table S7: Clinical profile of Proteomics dataset related to Figure 5A and Figure S5B.

Table S8: Protein to gene ratio of hallmark gene sets for proteomic analysis related to Figure 5E,

Table S9: Ligand-receptor output for NE, hybrid-NE and non-NE ecosystems using ICELLNET algorithm related to Figure 6A, 6B and Figure S6B.

Table S10: Differentially expressed Reactome pathways noted between non-NE vs NE/hybrid-NE TME segments related to Figure 6C.

Extended Data 1: Quantile 3 normalized processed data matrix for Spatial transcriptomics data set with metadata related to Figure 1, 2, 3, 4 and see STAR Methods.

Extended Data 2: Processed WGS data tumor mutation and copy number data related to Figure 1 and see STAR Methods.

Extended Data 3: Processed mass spectrometry proteomics data with metadata related to Figure 5 and see STAR Methods.

Extended Data 4: Processed bulk RNA-seq data for tumors used for proteomics analysis with metadata related to Figure 5 and see STAR Methods.

Extended Data 5: Processed bulk RNA-seq data of DMS 273 cell lines treated with control and Erdafitinib (33.33 micromolar concentration) with metadata related to Figure 6 and see STAR Methods.

Extended Data 6: Low magnification images of tumors profiled using spatial transcriptomics (GeoMx NanoString approach) with red circles highlighting the precise capture regions related to Figure 1 and see STAR Methods.

Extended Data 7: Low magnification images of tumors profiled used for mass spectrometry proteomics profiling with pre-LMD(left), post-LMD for tumor areas (center) and post-LMD for TME areas (right) included for each tumor related to Figure 5 and see STAR Methods.

---

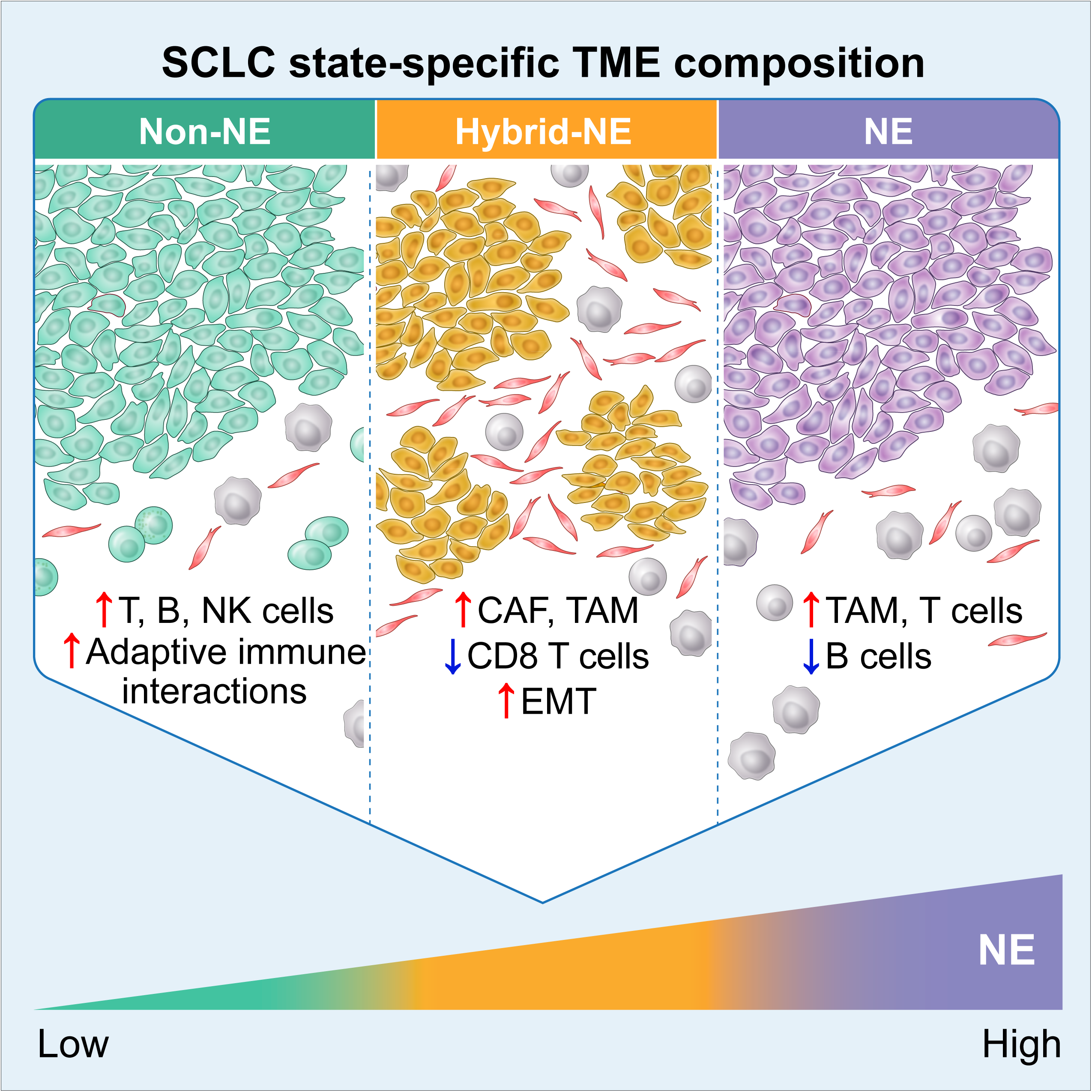

