## Supplemental file for "Microenvironment Shapes Small Cell Lung Cancer Neuroendocrine States and Presents Therapeutic Opportunities"

**Figure S1**

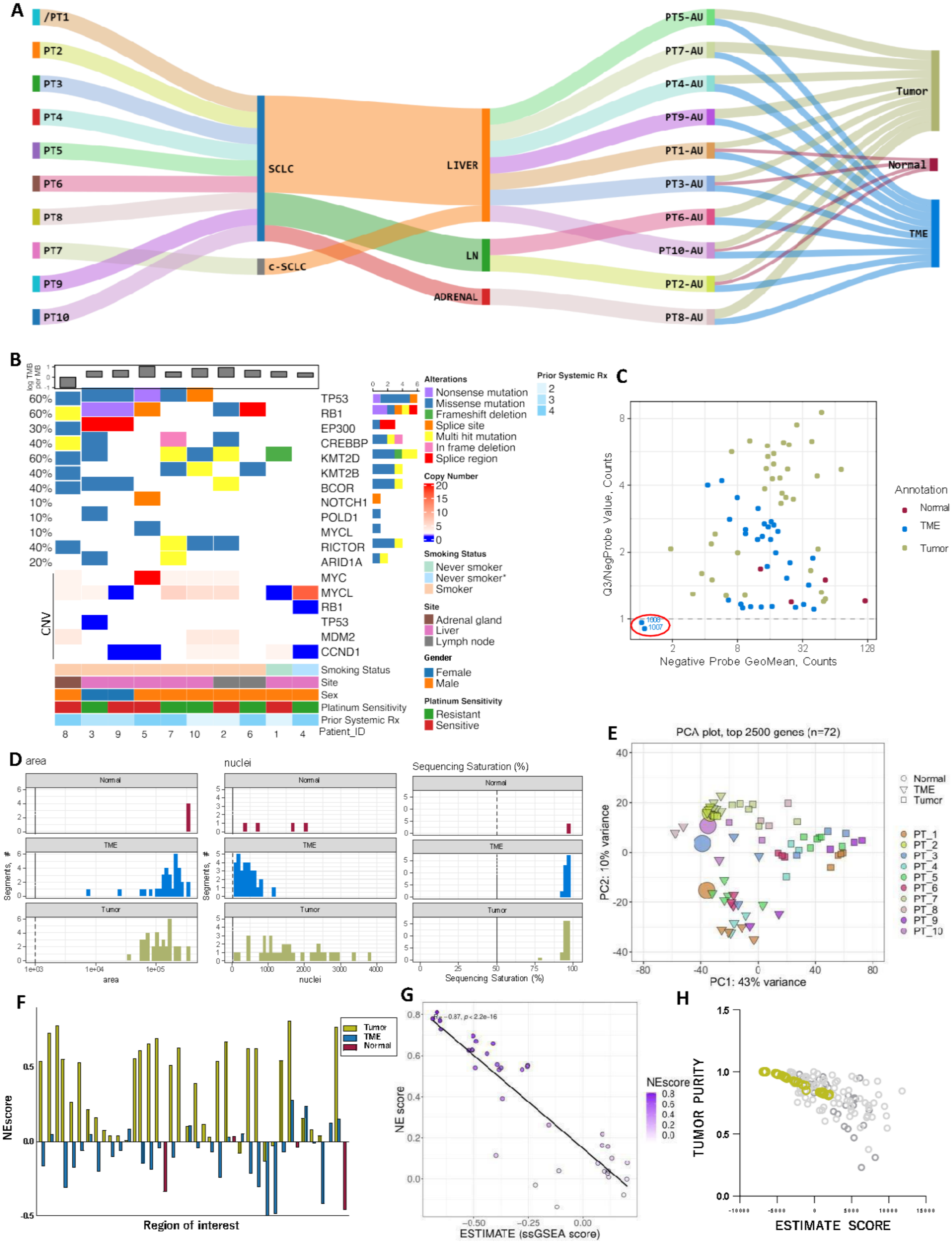
A
PT1
PT2
PT3
PT4
PT5
PT6
PT8
PT7
PT9
PT10
SCLC
c-SCLC
LIVER
LN
ADRENAL
PT5-AU
PT7-AU
PT4-AU
PT9-AU
PT1-AU
PT3-AU
PT6-AU
PT10-AU
PT2-AU
PT8-AU
Tumor
Normal
TME
B
log TMB per MB
TP53
RB1
EP300
CREBBP
KMT2D
KMT2B
BCOR
NOTCH1
POLD1
MYCL
RICTOR
ARID1A
MYC
MYCL
RB1
TP53
MDM2
CCND1
CNV
Smoking Status
Site
Sex
Platinum Sensitivity
Prior Systemic Rx
Patient_ID
Alterations
Nonsense mutation
Missense mutation
Frameshift deletion
Splice site
Multi hit mutation
In frame deletion
Splice region
Copy Number
Smoking Status
Never smoker
Never smoker*
Smoker
Site
Adrenal gland
Liver
Lymph node
Gender
Female
Male
Platinum Sensitivity
Resistant
Sensitive
Prior Systemic Rx
C
Q3/NegProbe Value, Counts
Negative Probe GeoMean, Counts
Annotation
Normal
TME
Tumor
D
area
nuclei
Sequencing Saturation (%)
Segments, #
E
PCA plot, top 2500 genes (n=72)
PC1: 43% variance
PC2: 10% variance
Normal
TME
Tumor
PT_1
PT_2
PT_3
PT_4
PT_5
PT_6
PT_7
PT_8
PT_9
PT_10
F
NEscore
Region of interest
Tumor
TME
Normal
G
NE score
ESTIMATE (ssGSEA score)
NEscore
H
TUMOR PURITY
ESTIMATE SCORE

**Figure S1: Spatial transcriptomics and whole genome sequencing profiling of metastatic and relapsed SCLC samples, corresponding to Figure 1**

A. Overall workflow of ST approach for spatially profiled tumors (n=10).

B. Mutational (above) and CNV (below) landscape of SCLC tumors (n=10) (whole genome sequencing) profiled using ST.

C. Quality metric of data assessed using mean Q3 value (of all 18,776 genes) to the negative probe Q3 value (y-axis) indicating TME segments 1007 and 1008 as outliers with relatively low-quality data (removed from subsequent analyses).

D. Bar-plots showing segment area of capture (left), number of nuclei (center), and sequencing saturation (right) for each tumor, TME and normal segments. Color code as Fig. S1C.

E. PCA plot showing PC1 vs PC2 like Fig 1B, additionally highlighting the precise location of normal (n=4), tumor(n=36) and TME(n=30) segments and colored by each patient origin. Overall normal segments clustered close to patient matched TME segments as opposed to tumor segments.

F. Bar plot showing NE score (ssGSEA) for each tumor (n=36), TME (n=30) and normal segment (n=4) profiled for each region.

G. Correlation between tumor segment NE scores (n=36) and their stromal and immune scores[40]. (Spearman correlation coefficient, r= -0.87).

H. Scatter plot showing the correlation between the tumor purity score projections (derived from linear regression modeling of bulk RNA and WGS sequencing data[5,9] using ABSOLUTE approach[41]to calculate tumor purity estimates and the stromal and immune scores[40] for spatially profiled tumor segment. Color code as Fig. S1C.

Abbreviations: ST- spatial transcriptomics; c-SCLC, combined small cell lung carcinoma; AU, autopsy; LN, Lymph node; CNV, copy number variations; Q3, 3rd Quantile normalized count, TME- tumor microenvironment, PC- principal component, PCA- principal component analysis, NE- neuroendocrine, Rx, treatment; ssGSEA, single sample gene set enrichment analysis; *non-smoker but very heavy exposure to asbestos.

### Figure S2

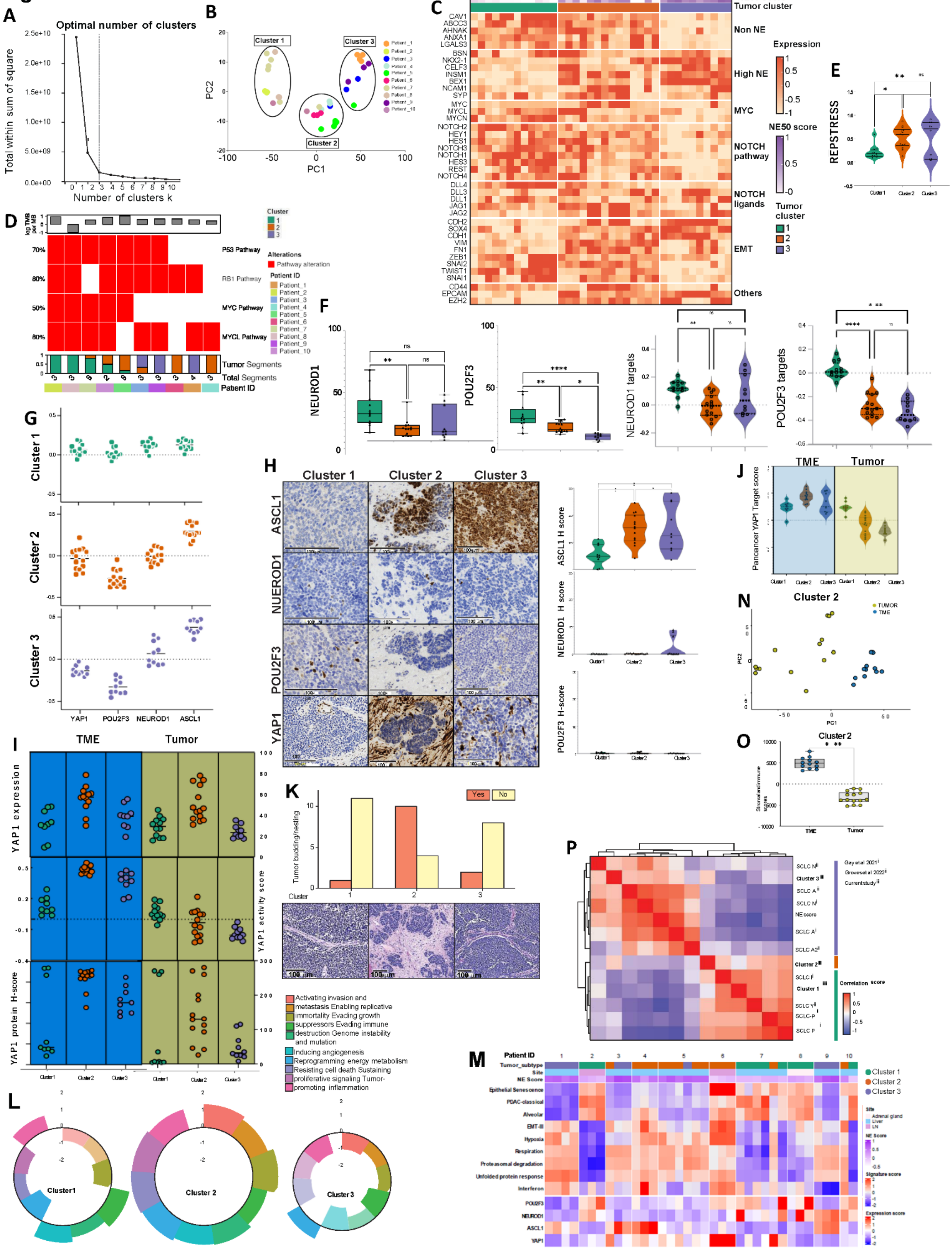

A
Optimal number of clusters
Total within sum of square
Number of clusters k
B
Cluster 1
Cluster 2
Cluster 3
PC1
PC2
C
NE50 score
Tumor cluster
Non NE
High NE
MYC
NOTCH pathway
NOTCH ligands
EMT
Others
Expression
Tumor cluster
D
P53 Pathway
RB1 Pathway
MYC Pathway
MYCL Pathway
Tumor Segments
Total Segments
Patient ID
Alterations
Pathway alteration
E
REPSTRESS
F
NEUROD1
POU2F3
NEUROD1 targets
POU2F3 targets
G
Cluster 1
Cluster 2
Cluster 3
YAP1
POU2F3
NEUROD1
ASCL1
H
ASCL1
NUEROD1
POU2F3
YAP1
ASCL1 H score
NEUROD1 H score
POU2F3 H-score
I
TME
Tumor
YAP1 expression
YAP1 activity score
YAP1 protein H-score
J
Pancancer YAP1 Target score
K
Tumor budding/nesting
Cluster
Yes
No
L
Cluster1
Cluster 2
Cluster 3
Activating invasion and metastasis Enabling replicative immortality Evading growth suppressors Evading immune destruction Genome instability and mutation
Inducing angiogenesis
Reprogramming energy metabolism
Resisting cell death Sustaining proliferative signaling Tumor-promoting inflammation
M
Patient ID
Tumor_subtype
Site
NE Score
Epithelial Senescence
PDAC-classical
Alveolar
EMT-III
Hypoxia
Respiration
Proteasomal degradation
Unfolded protein response
Interferon
POU2F3
NEUROD1
ASCL1
YAP1
Cluster 1
Cluster 2
Cluster 3
N
Cluster 2
TUMOR
TME
O
Cluster 2
TME
Tumor
P
Correlation score

**Figure S2 : Intratumoral spatial heterogeneity of SCLC neuroendocrine states and their spatial localization, related to Figure 2.**

A) Intra-cluster variation of the tumor segments (n=36) (as total within-sum of squares) (y-axis) plotted for each partitioning constant (k) for k=1-10. Relatively flattening of curve noted at k=3.

B) PCA as Fig 2A colored by patient origin to demonstrate intra-tumoral heterogeneity of tumor segments (n=36).

C) Heatmap showing distribution of SCLC related genes across the 3 clusters.

D) Oncoplot demonstrating spatially determined tumor cluster phenotype proportions in each (patient) tumor and presence of *RB1, TP53* pathway alterations (loss of function events) as well as *MYC* and *MYCL* alterations (amplification events) determined by bulk WGS.

E) Box- plots showing REPSTRESS[19] scores (ssGSEA derived) in three clusters.[#]

F) Gene expression counts (above, 3rd quantile normalized values) and target activity scores (below, ssGSEA) of *NEUROD1* (left) and *POU2F3* (right) in three tumor clusters. [#]

G) Cluster-wise landscape of SCLC lineage-defining transcription factor activity scores (ssGSEA derived).

H) Representative images at high power (40X magnification) showing protein expression of SCLC lineage-defining transcription factors (ASCL1,NEUROD1, POU2F3 and YAP1) across the tumor clusters. Scale bar at 100 µm. Quantification (right) showing expression of ASCL1, NEUROD1 and POU2F3 protein in 3 tumor clusters as H-scores (range from 0-300).[#]

I) Dot plots showing *YAP1* RNA expression (top, 3rd quantile normalized values), *YAP1* TF activity score (middle, ssGSEA derived) and IHC H-scores (bottom) across both TME and tumor segments demonstrating increased YAP1 activity in TME segments of cluster 2 followed by cluster 3.

J) Validation of distinct *YAP1* TF activity patterns across tumor and TME segments using independent pan-cancer *YAP1* signature[46].

K) Frequency of tumor budding in cluster 2 compared with cluster 1 and Cluster 3 SCLC. Representative images shown below. Scale bar at 100 µm.

L) Cancer hallmarks differentially enriched (GSEA derived) across the spatially profiled tumor clusters. The height of each bar shows NES.

M) Expression heatmap of transcript-defined cancer meta-programs[44] and SCLC lineage-defining transcription factors across different tumor segments from individual patient tumors (n=10).

N) PCA plot subsetted for only Cluster 2 regions colored by tumor (n=14) and TME (n=12) segments showing distinct clustering of tumor and TME (2500 most differentially expressed genes).

O) Stromal and immune (ESTIMATE) scores[40] showing negative enrichment of these scores in Cluster 2 tumor segments (n=14) as opposed to Cluster 2 TME segments (n=12).

P) Correlation matrix showing pairwise correlation for different SCLC related signatures with signatures generated for spatially profiled tumor segment clusters for SCLC cell lines (n=52) (like Fig. 2I).

Abbreviations: ST- spatial transcriptomics; WGS- whole genome sequencing, ssGSEA- single sample gene set enrichment analysis, IHC- immunohistochemistry, SCLC- small cell lung cancer; REPSTRESS, Replication stress; TF, transcription factor; *statistical significance at $p<0.05$; **statistical significance at $p<0.001$; ***statistical significance at $p<0.001$; ****statistical significance at $p<0.0001$; [#] Tukey's- multiple comparison test; NES-normalized enrichment score

#### Figure S3

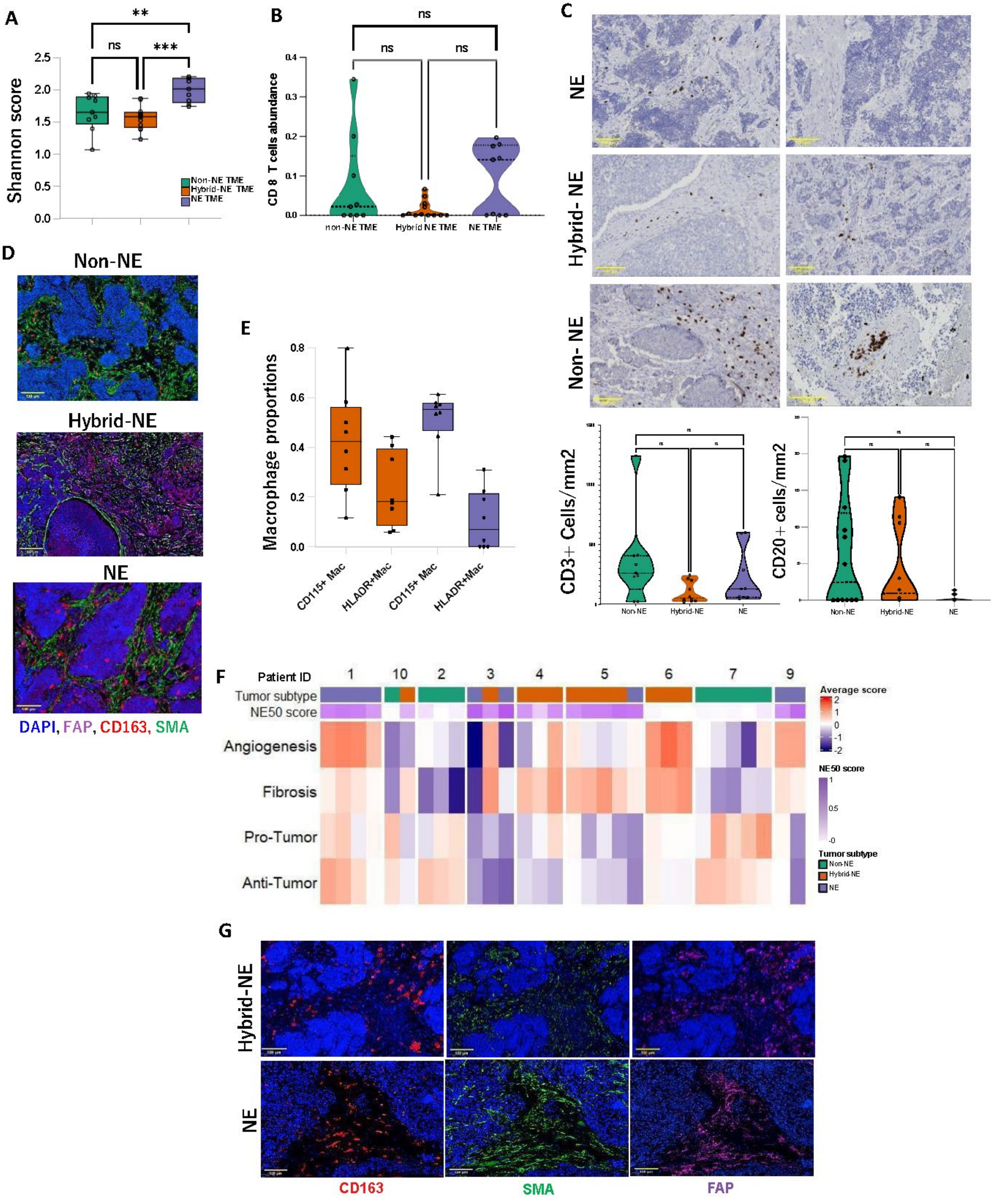

A
Shannon score
Non-NE TME
Hybrid-NE TME
NE TME
ns
**
***
B
CD 8 T cells abundance
non-NE TME
Hybrid NE TME
NE TME
ns
C
NE
Hybrid- NE
Non- NE
CD3+ Cells/mm2
CD20+ cells/mm2
Non-NE
Hybrid-NE
NE
D
Non-NE
Hybrid-NE
NE
DAPI, FAP, CD163, SMA
E
Macrophage proportions
CD115+ Mac
HLADR+Mac
F
Patient ID
Tumor subtype
NE50 score
Angiogenesis
Fibrosis
Pro-Tumor
Anti-Tumor
Average score
NE50 score
Tumor subtype
Non-NE
Hybrid-NE
NE
G
Hybrid-NE
NE
CD163
SMA
FAP

**Figure S3: SCLC TME characterization and association with spatially proximate SCLC tumor NE state related to Figure 3**

A) Shannon scores for CIBERSORT deconvoluted cell types for each TME subtype (n=30).[#]

B) CD8+ T cells abundance score (CIBERSORT) for each TME subtype (n=30). Color codes as Fig. S3A.[#]

C) Representative IHC images performed on sub-level sections (40x magnification) for CD3+ T cells (left) and CD20+ B cells (right) in different corresponding NE subtype tumor segments. Violin plots (below) demonstrating abundance of CD3+ T cells (left) and CD20+ B cells (right) [#]. Scale bar is set at 100µm.

D) Representative multispectral IF images corresponding to Fig. 3E (filters on for DAPI, CD163, SMA and FAP), non-NE (top), hybrid-NE (center), NE (bottom). Scale bar is set at 100µm.

E) Macrophage subtype proportions in NE and hybrid-NE TME.

F) Heatmap showing average expression scores of pan-cancer TME features[52] (reduced to 4 major TME features- fibrosis, angiogenesis, pro-tumor immune factors and anti-tumor immune factors) and clustered patient wise to demonstrate TME ITH. Color code of TME subtypes as Fig. S3A. Number labels on top indicate patient ID.

G) Single component (40x magnification) mIF images corresponding to Fig. 3G (patient#5 tumor) demonstrating individual staining of CD163 (left), SMA (center) and FAP (right). DAPI (blue) filter is on in all the images. Scale bar set at 100µm.

Abbreviations: TME, tumor microenvironment; IHC, immunohistochemistry; ITH, Intra-tumoral heterogeneity; DAPI,4′,6-diamidino-2-phenylindole; NE, neuroendocrine; FAP, Fibroblast activation protein- alpha; SMA, smooth muscle actin; Mac, macrophages; ns, statistically non-significant; **statistical significance at $p<0.001$; ***statistical significance at $p<0.001$; [#] Tukey's- multiple comparison test.

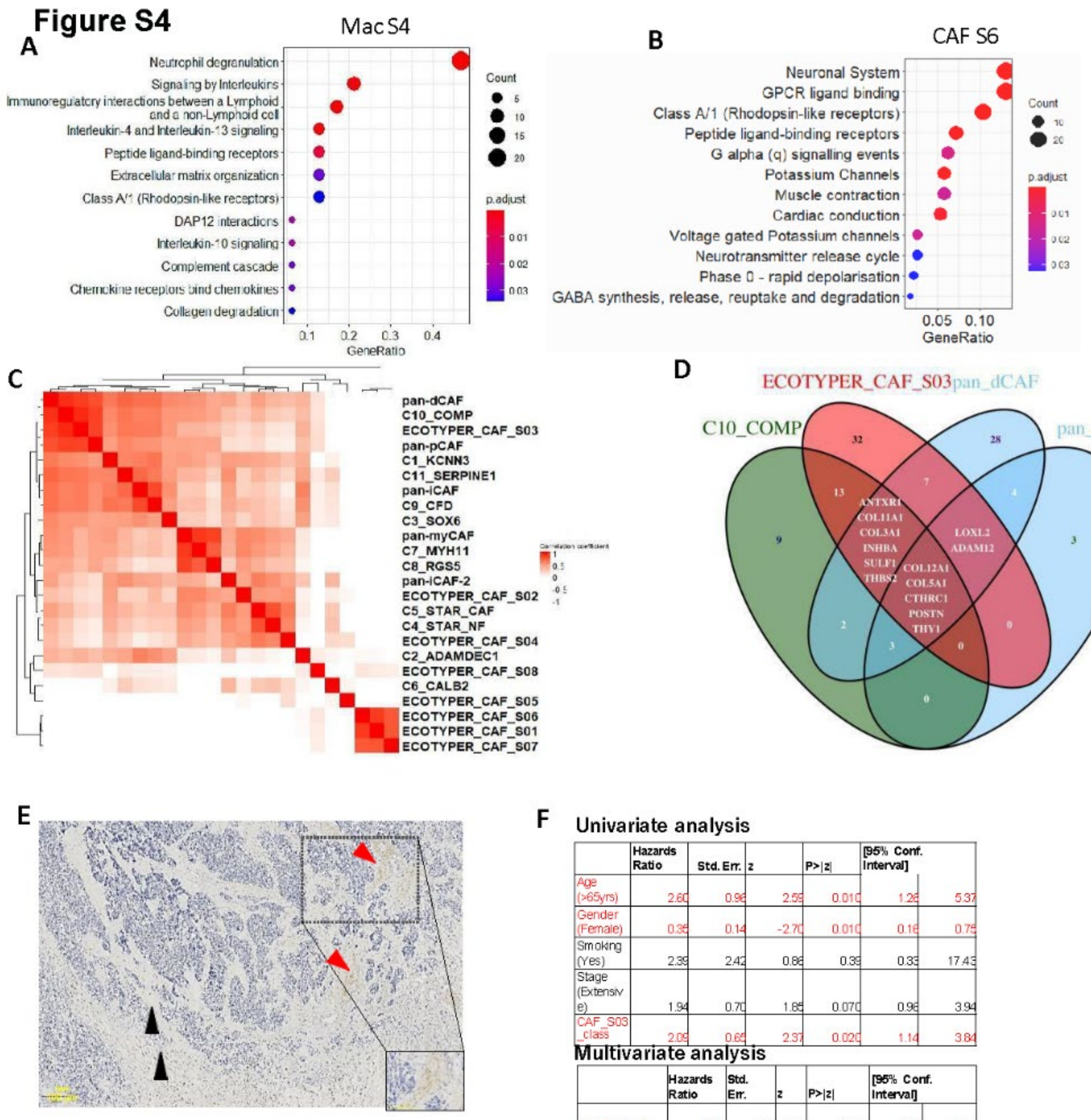


Univariate analysis

| | Hazards Ratio | Std. Err. | z | P>\|z\| | [95% Conf. Interval] | |
|---|---|---|---|---|---|---|
| Age (>65yrs) | 2.60 | 0.96 | 2.59 | 0.010 | 1.26 | 5.37 |
| Gender (Female) | 0.35 | 0.14 | -2.70 | 0.010 | 0.16 | 0.75 |
| Smoking (Yes) | 2.39 | 2.42 | 0.86 | 0.39 | 0.33 | 17.43 |
| Stage (Extensive) | 1.94 | 0.70 | 1.85 | 0.070 | 0.96 | 3.94 |
| CAF_S03_class | 2.09 | 0.65 | 2.37 | 0.020 | 1.14 | 3.84 |

Multivariate analysis

| | Hazards Ratio | Std. Err. | z | P>\|z\| | [95% Conf. Interval] | |
|---|---|---|---|---|---|---|
| Age (>65yrs) | 2.45 | 0.92 | 2.39 | 0.020 | 1.18 | 5.09 |
| Sex (Female) | 0.31 | 0.12 | -2.97 | 0.0030 | 0.14 | 0.67 |
| CAF_S03_class | 2.37 | 0.76 | 2.68 | 0.010 | 1.26 | 4.45 |

**Figure S4: Macrophage and CAF heterogeneity with their function defining biological heterogeneity in SCLC TME subtypes related to Figure 4.**

A) Programs enriched in Mac S4 cell state[57].

B) Programs enriched in CAF S6 cell state[57].

C) Pairwise-correlation plot of ssGSEA-derived enrichment scores of CAF signatures from different studies[57-59] in TME segments of spatially profiled tumors of the current study, like Fig 4C.

D) Common and distinct genes of different CAF types clustering together with ECOTYPER CAF S3[57].

E) Low power (20x magnification) IHC image of TEM8 (ANTXR1) in patient #10 tumor with inset showing positive TEM8 staining in TME of hybrid-NE segment (red arrows) same area corresponding to Fig. 2H. Negative staining in areas corresponding to non-NE TME segments (black arrows). Scale bar set at 100µm.

F) Univariate (above) and multivariate (below) survival analysis of bulk transcriptome dataset in SCLC with available survival data[2] considering CAF S03 high and low class (see methods). Cox proportional hazard algorithm used for survival analyses.

Abbreviations: Mac, macrophage; Mac S4- Macrophage cell state 4; CAF, cancer associated fibroblasts, Endo S2, Endothelial cell state 2; ssGSEA- single sample gene set enrichment analysis, TME- tumor microenvironment, SCNC- small cell neuroendocrine carcinoma, NE- neuroendocrine, IHC- immunohistochemistry, TEM8- tumor endothelial marker 8, std. error- standard error, **** statistical significance p<0.0001.

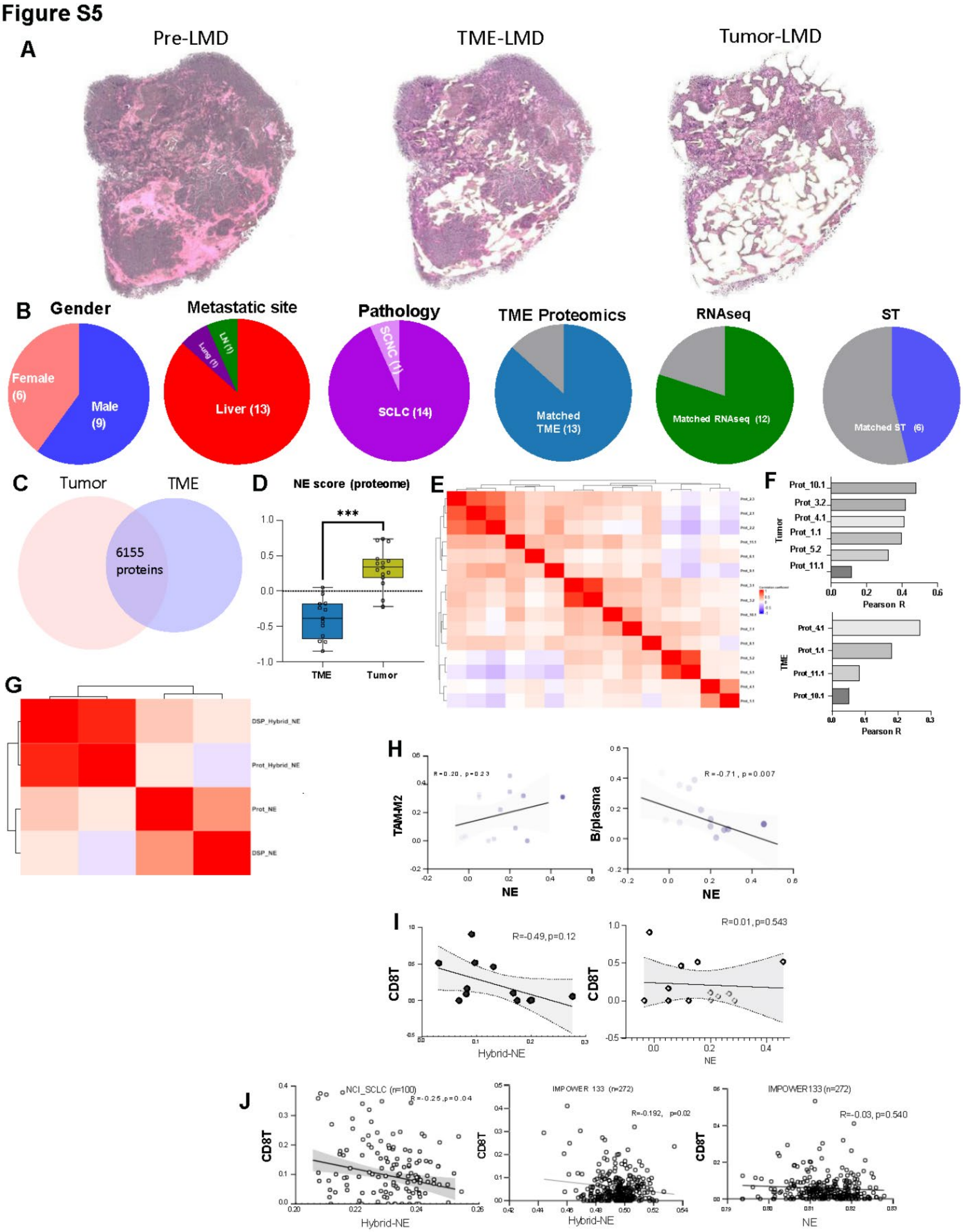
Figure S5
A
Pre-LMD
TME-LMD
Tumor-LMD
B
Gender
Female (6)
Male (9)
Metastatic site
Liver (13)
Lung (1)
LN (1)
Pathology
SCLC (14)
SCNC (1)
TME Proteomics
Matched TME (13)
RNAseq
Matched RNAseq (12)
ST
Matched ST (6)
C
Tumor
TME
6155 proteins
D
NE score (proteome)
***
TME
Tumor
E
F
Tumor
TME
Pearson R
Prot_10.1
Prot_3.2
Prot_4.1
Prot_1.1
Prot_5.2
Prot_11.1
G
DSP_Hybrid_NE
Prot_Hybrid_NE
Prot_NE
DSP_NE
H
TAM-M2
B/plasma
NE
R=-0.71, p=0.007
I
CD8T
R=-0.49, p=0.12
R=0.01, p=0.543
Hybrid-NE
NE
J
NCI_SCLC (n=100)
R=-0.25, p=0.04
IMPOWER 133 (n=272)
R=-0.192, p=0.02
IMPOWER133 (n=272)
R=-0.03, p=0.540
CD8T
Hybrid-NE
NE

**<u>Figure S5:</u> Proteomic profiling of relapsed and metastatic SCLC rapid autopsy tumors and tumor heterogeneity-linked reprogramming of SCLC TME related to Figure 5.**

A) Low power (4X), H&E image of a representative tumor tissue (prot#7.1) subjected to LMD for proteomics processing with pre-LMD (left), post-TME LMD (middle) and post-Tumor LMD (right).

B) Clinical and experimental distribution of proteomics profiled tumors (n=15) from tumors collected during rapid autopsy.

C) Venn diagram showing proteomic capture landscape of tumor and TME proteins in our dataset with 6155 common proteins.

D) Proteomics derived NE signature score[7,9] between tumor and TME enriched regions.[#]

E) Pairwise correlation of 1000 proteins with the highest variance in SCLC tumor proteome.

F) Transcript to protein correlation data for tumors with both proteome and spatial transcriptomics data available (n=6 for tumor enriched regions, n=4 for TME enriched regions).

G) Tumor transcript-protein correlation for NE and hybrid NE subtypes. Pairwise correlation of matched tumor proteome and ST tumor segments derived NE and hybrid-NE signatures (n=6).

H) Correlation of tumor proteome NE signature (x-axis; ssGSEA) with TME proteome-derived TAM-M2 (left) and B/plasma cells (right) (CIBERSORT-derived proportions).

I) Correlation of tumor proteome NE (right) and Hybrid-NE (left) signature (ssGSEA) with TME proteome-derived CD8T signatures (CIBERSORT- derived).

J) Correlation of Hybrid-NE (left, middle) and NE signature (right) (ssGSEA) with CD8 T signatures (CIBERSORT-derived) in larger SCLC bulk-RNA sequencing dataset[9,16].

Abbreviations: LMD, laser capture microdissection; TME, tumor microenvironment; H&E, hematoxylin, and eosin; ssGSEA, single sample gene set enrichment analysis; ST, Spatial transcriptomics; DSP, digital spatial profiling-Spatial transcriptomics; CAF, cancer associated fibroblasts; ****statistical significance at $p<0.0001$, [#] student t-test.

### Figure S6

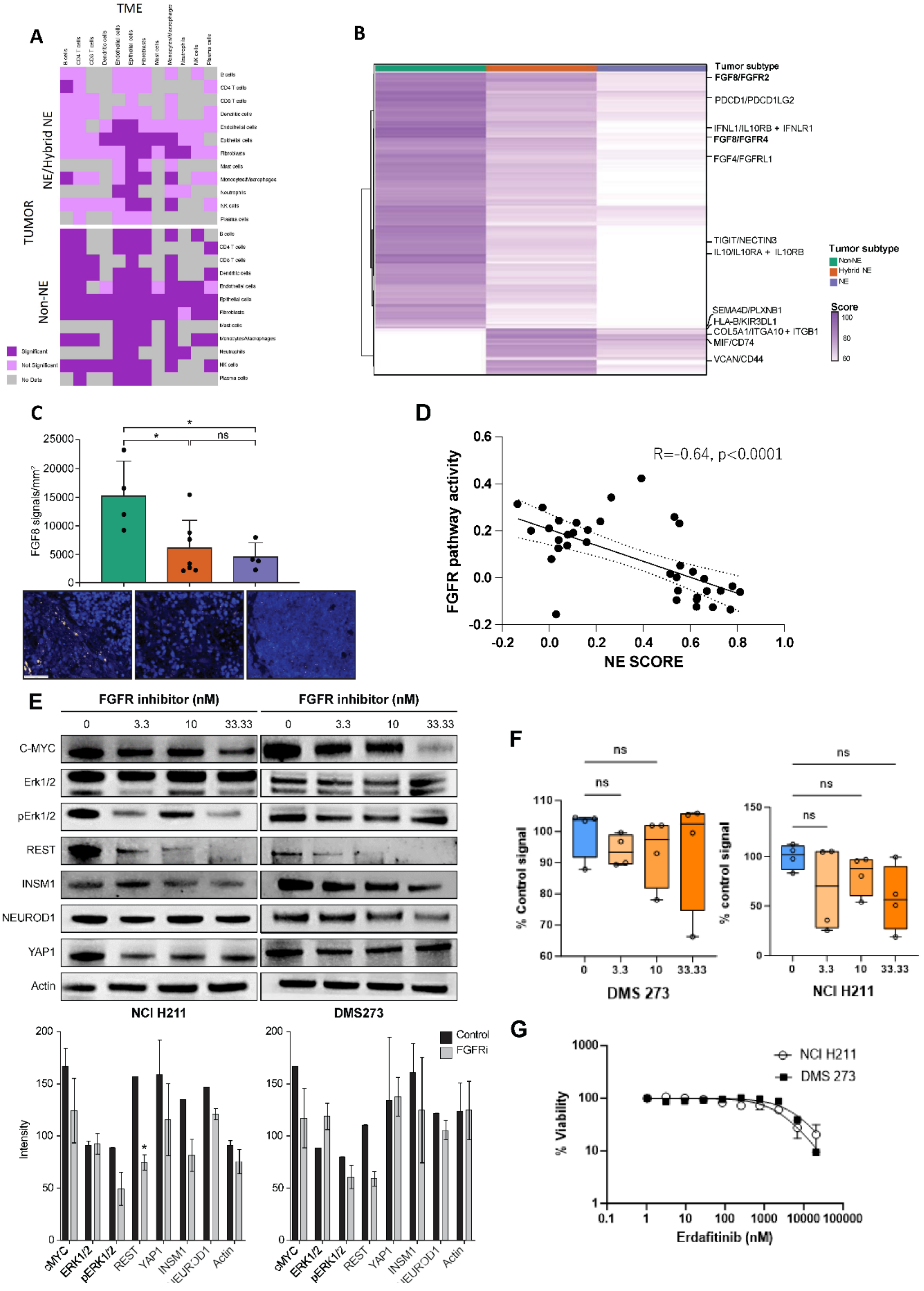

A
TME
TUMOR
NE/Hybrid NE
Non-NE
Significant
Not Significant
No Data
B
Tumor subtype
FGF8/FGFR2
PDCD1/PDCD1LG2
IFNL1/IL10RB + IFNLR1
FGF8/FGFR4
FGF4/FGFRL1
TIGIT/NECTIN3
IL10/IL10RA + IL10RB
SEMA4D/PLXNB1
HLA-B/KIR3DL1
COL5A1/ITGA10 + ITGB1
MIF/CD74
VCAN/CD44
Non-NE
Hybrid NE
NE
Score
C
FGF8 signals/mm²
D
R=-0.64, p<0.0001
FGFR pathway activity
NE SCORE
E
FGFR inhibitor (nM)
C-MYC
Erk1/2
pErk1/2
REST
INSM1
NEUROD1
YAP1
Actin
NCI H211
DMS273
Control
FGFRi
Intensity
F
% Control signal
% control signal
DMS 273
NCI H211
G
NCI H211
DMS 273
% Viability
Erdafitinib (nM)

**<u>Figure S6</u>: Modulation of SCLC tumor NE state by abrogation of FGF-FGFR signaling related to Figure 6.**

A) Heatmap of global interactions between Tumor (receiver) and TME (sender) using an orthogonal cell-cell interaction approach (Cellphone DB)[66] confirming increased and varied significant interactions between tumor and TME regions in non-NE ecosystems compared to NE/Hybrid-NE ecosystems.

B) Heatmap of ligand-receptor interaction pairs using iCELLNET5 showing most differentially enriched interactions (TME→ tumor). Clinically relevant and potentially targetable interactions are highlighted. FGF8 related interactions in bold.

C) *FGF8* RNA ISH signals across TME subtypes. Representative images (high power 40x, magnification) showing *FGF8* (yellow) in non-NE TME (left), hybrid-NE TME (middle) and NE TME (right)[#]. Nuclei are blue (DAPI). Scale bar set at 50µm.

D) *FGFR* activity scores (ssGSEA) in spatially resolved tumor segments transcriptomic (n=36) data.

E) FGF signaling intermediates, NE, and non-NE proteins in NCI-H211 and DMS-273 following treatment with erdafitinib at varying concentrations. (Below) Quantification of western blots of FGF signaling intermediates, NE, and non-NE proteins in NCI-H211 and DMS-273 after treatment with erdafitinib at varying concentrations.

F) Caspase-8 activation assay showing no significant increase in apoptosis at day 5 after erdafitinib treatment in DMS 273(left) and NCI H211 (right) SCLC cell lines.[&]

G) Cell titer glow viability assay showing no decrease in cell viability at erdafitinib concentrations used in this experiment.

Abbreviations: ssGSEA- single sample gene set enrichment analysis, FGF- fibroblast growth factor, * statistical significance at $p<0.05$; R= spearman correlation co-efficient; [#] student t-test; [&] Tukey's multiple comparison test.

Extended Data 6

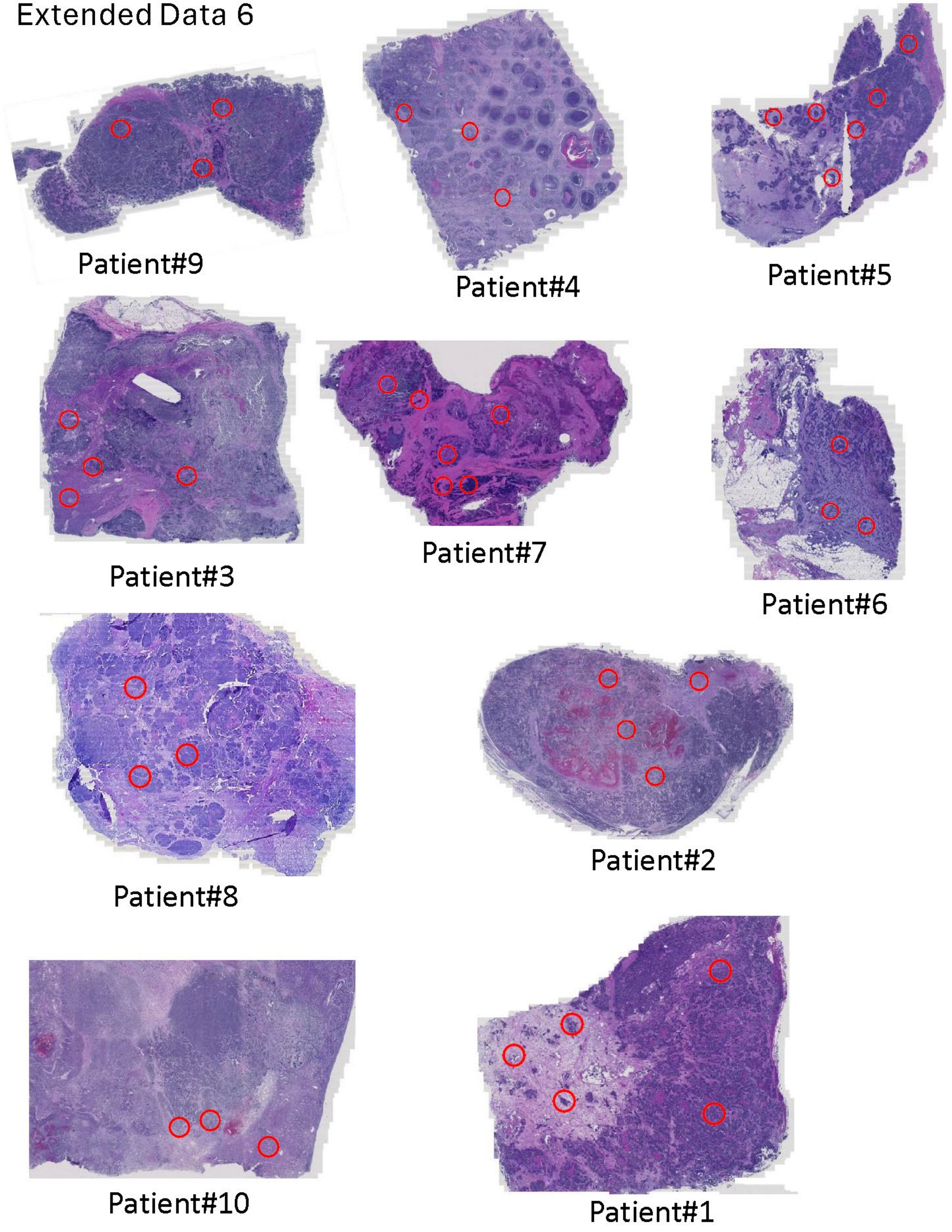

### Extended Data 7

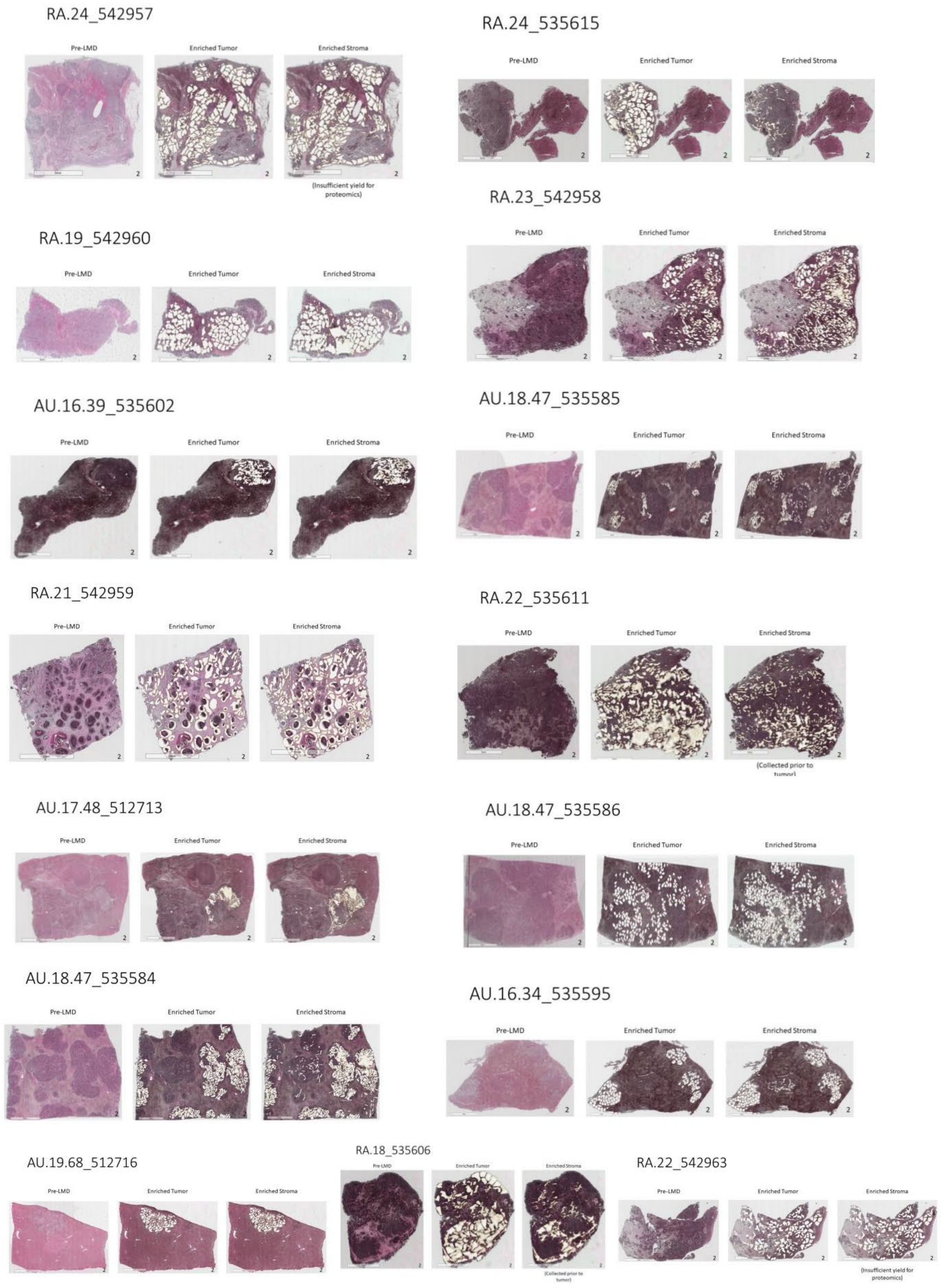
